# Human transmembrane protein 68 links triacylglycerol synthesis to membrane lipid homeostasis

**DOI:** 10.1101/2024.04.14.589399

**Authors:** Fansi Zeng, Christoph Heier, Qing Yu, Huimin Peng, Feifei Huang, Zheng Zhao, Pingan Chang

## Abstract

Transmembrane protein 68 (TMEM68) is a recently identified mammalian triacylglycerol (TAG) synthase with high expression in the brain. How TMEM68 regulates cellular lipid metabolism in concert with other enzymatic pathways remains poorly understood. In this study, we assessed TMEM68 function in neuro- and gliablastoma cells by combining genetic gain- and loss-of-function approaches with lipidomics. We found that TMEM68 promotes TAG synthesis and lipid droplet formation independently of canonical acyl-CoA:diacylglycerol acyltransferase (DGAT) 1 and 2 enzymes and contributes a discrete fraction of basal cellular TAG storage. Notably, TMEM68 overexpression increased storage lipids at the expense of membrane lipids leading to a profound reduction of ether-linked glycerophospholipids (GPLs). Moreover, altered TMEM68 expression levels were associated with diminished prevalence of polyunsaturated GPLs. We conclude that TMEM68, besides its TAG synthesis function, acts as a multifaceted regulator of membrane lipid composition and polyunsaturated fatty acid homeostasis.

## 1 Introduction

The synthesis of triacylglycerol (TAG) in mammals serves crucial roles in various physiological processes, such as intracellular storage of surplus energy, attenuation of lipotoxicity, intestinal dietary fat absorption, lactation, lipid transportation and signal transduction (*Shi and Cheng, 2009*). Mammalian TAG synthesis critically depends on glycerolipid acyltransferase enzymes that catalyze the sequential esterification of fatty acids (FAs) to a glycerol backbone using acyl-CoA as donor (*Cao et al., 2012*). Mammals engage two distinct biochemical pathways for TAG biosynthesis, (*Shi and Cheng, 2009; Wang et al., 2017*). The glycerol-3-phosphate (G3P) pathway, also known as the *de novo* synthesis pathway contributes the majority of TAG in most cells and tissues (*Wang et al., 2017*) whereas the monoacylglycerol (MAG) pathway operates in a restricted set of tissues including the small intestine, adipose tissue, and liver (*Yen et al., 2015; Stone, 2022*). The G3P pathway is initiated by glycerol-3-phosphate acyltransferase (GPAT) enzymes, which catalyze the formation of 1-acylglycerol-3-phosphate (also known as lysophosphatidic acid, LPA) from G3P and acyl-CoA (*Gimeno and Cao, 2008; Wendel et al., 2009*). Then, acylglycerophosphate acyltransferase (AGPAT) acylates LPA at the *sn*-2 position to produce phosphatidic acid (PA). PA is dephosphorylated by PA phosphatase (PAP) to diacylglycerol (DAG) (*Takeuchi and Reue, 2009*), which is finally acylated to TAG by acyl-CoA:diacylglycerol acyltransferase (DGAT) enzymes (*Shi and Cheng, 2009*). The MAG pathway is initiated by the acylation of MAG to DAG, which is catalyzed by MAG acyltransferase (MGAT) enzymes and thereafter converges with the G3P pathway at the DGAT step (*Shi and Cheng, 2009; Yen, et al., 2015; Stone, 2022*). TAG synthesis intersects with various pathways for glycerophospholipid (GPL) biosynthesis as PA and DAG are precursors for phosphatidylcholine (PC), phosphatidylethanolamine (PE), phosphatidylserine (PS), phosphatidylinositol (PI), phosphatidylglycerol (PG) and cardiolipin (CL) (*Farese and Walther, 2022; Lee and Ridgway, 2020*).

In mammals, two DGAT isoenzymes termed DGAT1 and DGAT2 account for the majority of TAG synthesis in most cell types and tissues (*Harris et al., 2011; Li et al., 2015; Roe et al., 2018*). Although DGAT1 and DGAT2 can operate in redundancy they belong to structurally unrelated protein families and differ in subcellular localization, membrane topology, and substrate specificity (*Chen et al., 2022*). Accordingly, several specific cellular and physiological functions have been ascribed to DGAT1 and DGAT2, respectively, including non-redundant roles in FA detoxification, lipid droplet (LD) subset formation, and maintenance of intestinal and epidermal integrity (*Chen et al., 2022; Stone, 2022*). Homologs of DGAT1 and DGAT2 enzymes are found in all major eukaryotic kingdoms. However, plants and fungi employ also a well-described acyl-CoA independent pathway as a major route of TAG synthesis, in which FAs are transferred from phospholipids to DAG via the enzyme phospholipid:diacylglycerol acyltransferase (PDAT) (*Xu, et al., 2018; Sah, et al., 2023*). While mammalian homologs of plant and fungal PDATs exist (e.g. Lecithin:cholesterol acyltransferase and Lecithin:retinol acyltransferase), these enzymes prefer acyl acceptors other than DAG and are therefore not considered to contribute to mammalian TAG synthesis (*Sears et al., 2016; Falarz et al., 2020*). Thus, at present DGAT1 and DGAT2 are regarded as the major if not sole enzymes responsible for TAG synthesis and LD formation in mammalian cells. Nevertheless, residual cellular TAG has been observed after combined genetic ablation or pharmacological inhibition of DGAT1 and DGAT2 arguing for the presence of additional enzymes in mammalian TAG synthesis (*Harris et al., 2011; Chitraju et al., 2023*).

One candidate enzyme recently implicated in mammalian TAG synthesis is transmembrane protein 68 (TMEM68), a poorly characterized acyltransferase with high expression in the brain (*Chang et al., 2017*). TMEM68 shares the pfam01553 acyltransferase domain with several GPAT and AGPAT isoenzymes (*Shan et al., 2010*). Of note, however, TMEM68 contains only motif I of the pfam01553 domain, but unlike GPAT/AGPAT lacks the conserved substrate binding motifs II and III (*Chang et al., 2017; Valentine et al., 2022; Yamashita et al., 2014*) suggesting that TMEM68 differs in substrate specificity from GPAT/AGPAT enzymes. We have previously shown that TMEM68 overexpression in mammalian cells promotes TAG biosynthesis and LD accumulation in a conserved active sites-dependent manner (*Wang et al., 2023*). The ectopic expression of TMEM68 in HEK293 cells increased TAG storage along with cellular MGAT and DGAT activities arguing for a MGAT/DGAT-like function of this enzyme (*Wang et al., 2023*). A recent study confirmed the function of TMEM68 in TAG synthesis yet attributed it to a possible PDAT rather than MGAT/DGAT activity of the enzyme (*McLelland et al., 2023*). Considering the high expression of TMEM68 in the brain, in the current study we set out to assess the broader role of TMEM68 in lipid metabolism of neuro- and gliablastoma cell models. Using loss- and gain-of-function approaches we found that TMEM68 autonomously promotes TAG and LD formation independent of DGAT1/2 and in addition regulates molecular composition of cellular GPLs.

## Results

### TMEM68 regulates TAG levels in neuroblastoma and glioblastoma cells

Our previous study revealed that TMEM68 overexpression promotes TAG accumulation in human embryonic kidney HEK293 and breast cancer MCF-7 cells (*Wang et al., 2023*). The mRNA levels of *TMEM68* in the mouse brain are higher than those in other tissues suggesting a particular role of the enzyme in cell types of the central nervous system (*Chang et al., 2017*). In line with brain-enriched expression of *TMEM68*, qPCR analysis revealed higher *TMEM68* mRNA levels in human neuroblastoma SK-N-SH and glioblastoma U251 cells compared to HEK293 cells (***Figure 1-figure supplement 1***). To assess the possible function(s) of TMEM68 in glial/neuronal lipid metabolism, we first generated cell lines stably expressing TMEM68 or green fluorescence protein (GFP), respectively, and analyzed cellular TAG levels. As shown in ***Figure 1A***, mRNA levels of *TMEM68* in TMEM68-expressing SK-N-SH (SK/TMEM68) and U251 (U251/TMEM68) cells were more than 100-fold higher compared with GFP-expressing SK/GFP and U251/GFP control cells. TMEM68 protein levels in SK/TMEM68 cells were also substantially increased compared to SK/GFP cells (***Figure 1B***). In line with previous findings (*Wang et al., 2023*), TMEM68-overexpression was associated with increased TAG levels in SK/TMEM68 compared with SK/GFP cells both in the absence (6.1-fold) and presence of exogenous oleic acid (OA, 3.9-fold) (***Figure 1C***). Similarly, overexpression of TMEM68 elevated TAG levels in U251 cells in the absence (1.7 fold) or presence of OA (1.5-fold) relative to GFP-expressing cells (***Figure 1D***). To further investigate the function of TMEM68 in cellular TAG metabolism, we depleted endogenous TMEM68 in human SK-N-SH and U251 cells by CRISPR/Cas9-mediated gene editing and RNA interference (RNAi), respectively. Stable expression of two gRNA constructs targeting TMEM68 (TMEM68 knockout (KO) 1 and KO2) decreased the *TMEM68* mRNA concentration by 71% and 97% respectively, as compared with unmodified SK-N-SH control cells (***Figure 1E***). This resulted in undetectable TMEM68 protein levels in SK/TMEM68 KO2 cells (***Figure 1F***). Among the three shRNA constructs targeting TMEM68 (shTMEM68) only shTMEM68-1 robustly downregulated *TMEM68* mRNA levels in U251 cells by more than 70% (***Figure 1E***, ***Figure 1-figure supplement 2***). Thus, SK/TMEM68 KO2 and U251/shTMEM68-1 cells were used in the next experiments and referred to as SK/TMEM68 KO and U251/shTMEM68 respectively. Reduced TMEM68 expression led to a significant 25-31% decrease in total TAG levels in SK-N-SH and U251 cells compared to controls both, in the absence and presence of exogenous OA (***Figure 1G*, *H***). Since TAG is preferentially stored in intracellular LDs we stained those compartments with Nile Red and imaged them by confocal fluorescence microscopy. As shown in ***Figure 1I***, LDs were apparently larger and more abundant in SK/TMEM68 cells than in SK/GFP cells. In contrast, we observed fewer LDs in SK/TMEM68 KO cells relative to SK-N-SH control cells. Taken together, these data identify TMEM68 as a positive regulator of TAG and LD storage in neuro- and glioblastoma cells.

**Figure 1.**
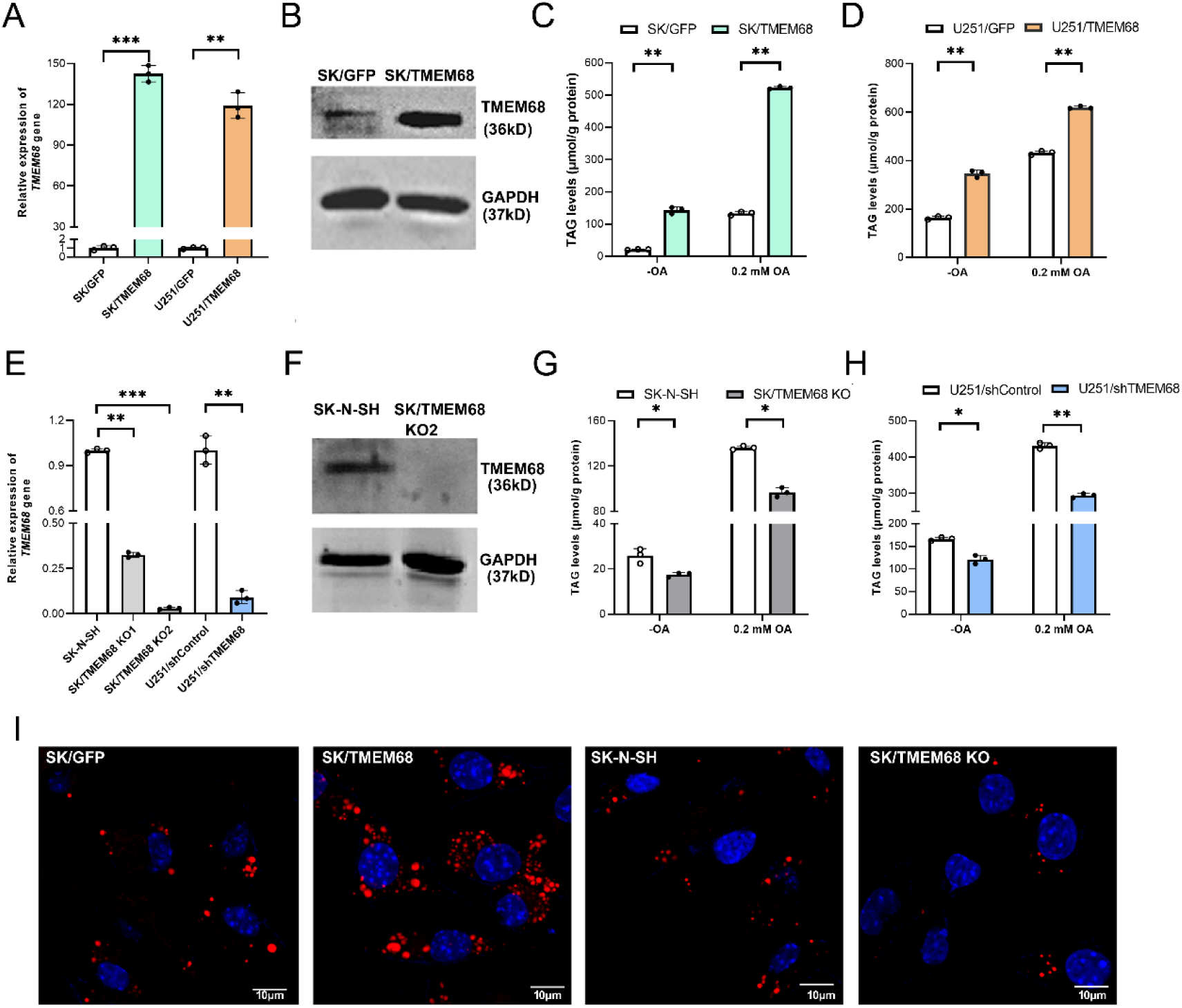
TMEM68 promotes cellular TAG and LD formation. (**A**, **E**) *TMEM68* mRNA levels in (**A**) TMEM68-expressing SK/TMEM68 and U251/TMEM68 cells, as well as in GFP-expressing SK/GFP and U251/GFP control cells; (**E**) in TMEM68-deficient TMEM68 KO (SK/TMEM68 KO) and TMEM68 knockdown (U251/shTMEM68) cells, as well as in SK-N-SH and U251/shControl control cells. (**B**, **F**) TMEM68 protein expression was detected by immunoblotting in (**B**) SK/TMEM68 and SK/GFP control cells; (**F**) SK/TMEM68 KO2 and SK-N-SH control cells using an anti-TMEM68 antibody. GAPDH protein expression was detected as loading control. (**C**, **D**, **G**, **H**) TAG levels were determined in the presence or absence (−OA) of 0.2 mM OA in (**C**) SK/TMEM68 and SK/GFP cells, (**D**) U251/TMEM68 and U25/GFP cells, (**G**) SK/TMEM68 KO and SK-N-SH cells as well as in (**H**) U251/shTMEM68 and U25/shControl cells. (**I**) LDs and nuclei were stained with HSC LipidTOX^TM^ Deep Red and DAPI respectively. Images were captured by confocal fluorescence microscopy. Figures are representative of at least three experiments. Data is presented as means ± SD. Asterisk indicates *P-*values: * *P* < 0.05, ** *P* < 0.01, *** *P* < 0.001, *n* = 3.

### TMEM68 expression controls cellular neutral lipid and FA levels

To further investigate how TMEM68 affects intracellular lipid metabolism, we compared lipids of SK/TMEM68, SK/TMEM68 KO, and control cells by quantitative untargeted LC/MS. A total of 555 lipids, including fatty acyls, glycerolipids, GPLs, sphingolipids and sterols were detected. TAG species accounted for nearly 30% of the total lipids detected (***Figure 2-figure supplement 1***). Consistent with our biochemical assays, total TAG was significantly increased 5.8-fold in SK/TMEM68 as compared to SK/GFP control cells (***Figure 2A***). MAG and DAG, the precursors of TAG in the MAG pathway, were also elevated upon TMEM68 overexpression by 3.2- and 2.2-fold, respectively (***Figure 2A*)**. Moreover, SK/TMEM68 cells exhibited 3.5-fold increased levels of cholesterol ester (CE), which like TAG is typically stored in LDs (***Figure 2A*)**. Conversely, TMEM68-deficiency reduced total DAG and TAG levels by 36% and 31%, respectively, although total MAG and CE levels were not significantly altered (***Figure 2D***). By analyzing DAG species we found that TMEM68 overexpression broadly increased the levels of most DAG species composed of saturated fatty acids (SFAs) and monounsaturated fatty acids (MUFAs) yet decreased the levels of specific DAG species containing 18:2 or 20:4 polyunsaturated fatty acids (PUFAs). As a consequence, combined DAG species with 0 and 1 double bond were increased, while DAG species with 4 double bonds were decreased in SK/TMEM68 cells compared to controls (***Figure 2B***). TMEM68-deficiency reduced the abundance of ∼50% of the detected DAG species without an apparent bias regarding FA saturation (***Figure 2E***). The analysis of TAG species revealed that TMEM68 overexpression preferentially increased saturated and monounsaturated TAGs (19.1- and 11.5- fold compared to controls) as compared to TAG with higher saturation degree (1.0-7.2 fold compared to controls) (***Figure 2C*, *Figure 2-figure supplement 2A***). Conversely, SK/TMEM68 KO cells exhibited a preferential decrease in unsaturated TAG with 5 or more double bonds (0.30-0.48 fold compared to controls) as compared to saturated or monounsaturated TAG (approximately 0.6-fold) (***Figure 2F*, *Figure 2-figure supplement 2B***).

**Figure 2.**
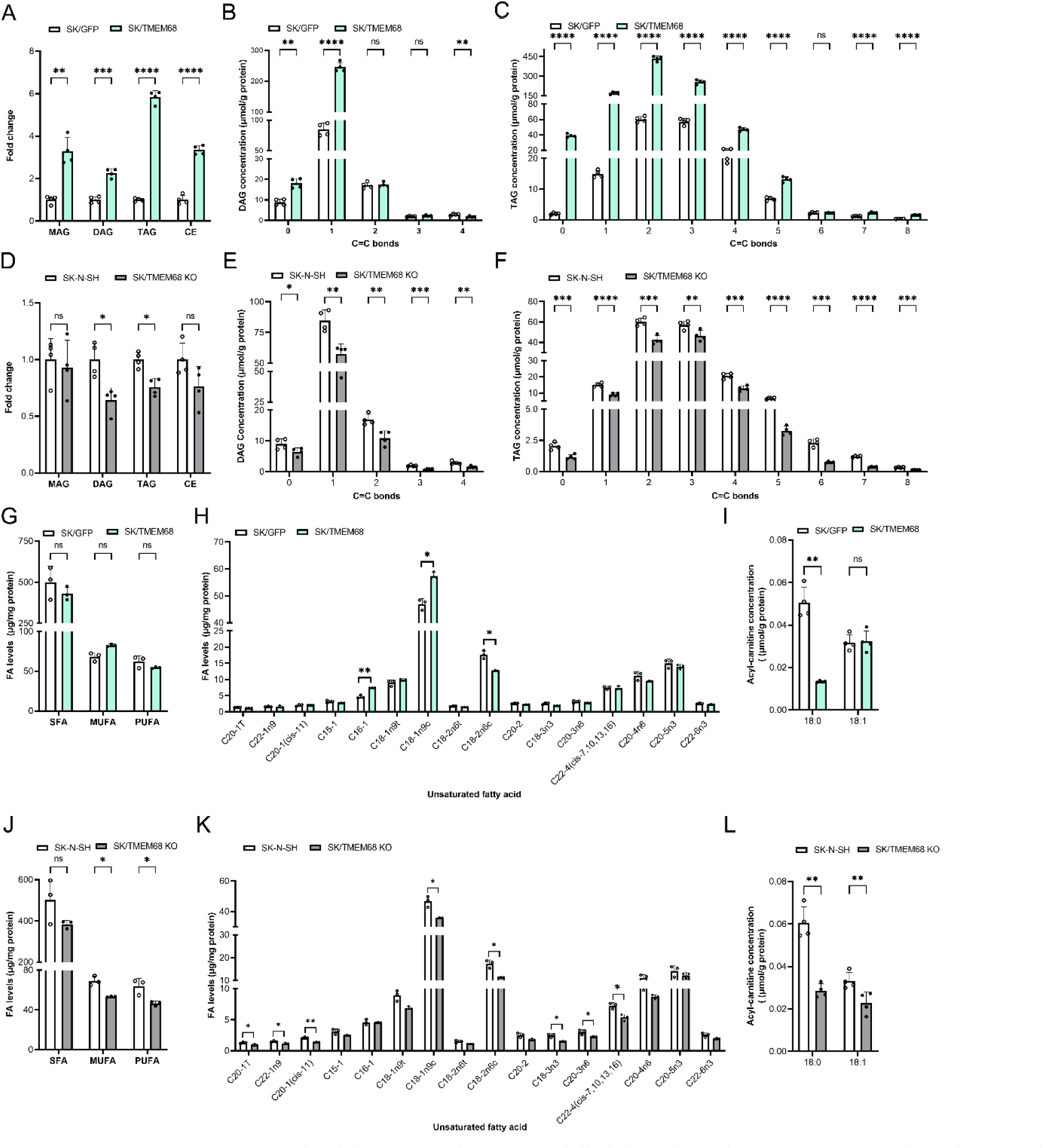
TMEM68 controlled intracellular neutral lipid and FA levels. (**A**, **D**) Abundance of total MAG, DAG, TAG and CE in (**A**) SK/TMEM68 and SK/GFP control cells; (**D**) SK/TMEM68 KO and SK-N-SH control cells. (**B**, **E**) Combined double bonds in DAG species of (**B**) SK/TMEM68 and SK/GFP cells; (**D**) SK/TMEM68 KO and SK-N-SH cells. (**C, F**) Combined double bonds in TAG species of (**C**) SK/TMEM68 and SK/GFP cells; (**F**) SK/TMEM68 KO and SK-N-SH cells. (**G, J**) Levels of total SFA, MUFA, and PUFA in (**G**) SK/TMEM68 and SK/GFP cells; (**J**) SK/TMEM68 KO and SK-N-SH cells. (**H, K**) Unsaturated FA levels in (**H**) SK/TMEM68 and SK/GFP cells; (**K**) SK/TMEM68 KO and SK-N-SH cells. (**I, L**) Levels of fatty acyl carnitines in (**I**) SK/TMEM68 and SK/GFP cells; (**L**) SK/TMEM68 KO and SK-N-SH cells. Data is presented as means ± SD. Statistically significant differences are marked with asterisks indicating *P*-values. * *P* < 0.05, ** *P* < 0.01, *** *P* < 0.001, **** *P* < 0.0001. ns, not significant. *n* = 4.

The distinct effects of TMEM68 on DAG and TAG species with different saturation degrees led us to investigate how TMEM68 expression affects non-esterified FAs. The targeted metabolomic analysis showed that TMEM68 overexpression significantly increased MUFAs, but not SFAs or PUFAs, whereas TMEM68-deficiency decreased MUFAs and PUFAs, but not SFAs (***Figure 2G*, *2J***). Palmitoleic acid (C16-1) and oleic acid (cis-9-octadecenoic acid, C18-1n9c) were the main FAs increased in SK/TMEM68 cells compared to SK/GFP control cells whereas linoleic acid (C18:2) levels were significantly decreased (***Figure 2H***). TMEM68-deficiency significantly reduced levels of eicosenoic acid (C20:1), oleic acid (C18:1), linoleic acid (C18:2), linolenic acid (C18:3), and docosatetraenoic acid (C22:4) as compared to controls (***Figure 2K***). The abundance of SFA species was unaffected by TMEM68 expression (***Figure 2-figure supplement 3***). However, among two forms of fatty acyl carnitines 18:0-carnitine content was decreased by TMEM68 overexpression and deficiency (***Figure 2I*, *2L***). These data together demonstrate that TMEM68 expression regulates neutral lipid metabolism beyond TAG by affecting abundance of acylglycerol intermediates, non-esterified FA, and acyl carnitines.

### TMEM68 controls TAG independently of DGAT1 and DGAT2

Given the broad impact of TMEM68 expression on cellular neutral lipids and FA, we next assessed mRNA expression of genes related to FA and TAG metabolism using RT-qPCR. TMEM68 overexpression significantly decreased mRNA levels of *DGAT1* but not *DGAT2* (***Figure 3A***). On the other hand, expression of *adipose triglyceride lipase* (*ATGL*), a key regulator of TAG mobilization, was unchanged in SK/TMEM68 cells as compared to controls (***Figure 3A***). Notably, expression of several genes involved in *de novo* FA synthesis, FA transport and the G3P pathway was elevated in SK/TMEM68 cells compared to controls, including mRNA levels of *fatty acid synthase* (*FASN*), *acetyl-CoA carboxylase α* (*ACACA*), *stearoyl-CoA desaturase 1* (*SCD1*), *fatty acid binding protein 4* (*FABP4*), *GPAT*3, *GPAT*4, *AGPAT*3, *AGPAT*4 and *LPIN*1 (***Figure 3A***). These analyses imply that increased FA *de novo* synthesis and upregulation of G3P pathway enzymes contribute to enhanced TAG deposition in SK/TMEM68 cells. In contrast, TMEM68-deficiency had little effect on lipid metabolic gene expression, the exceptions being an increase in *LPIN*1 mRNA concentration and a decrease in *FASN* and *FABP4* mRNA concentrations (***Figure 3B***).

**Figure 3.**
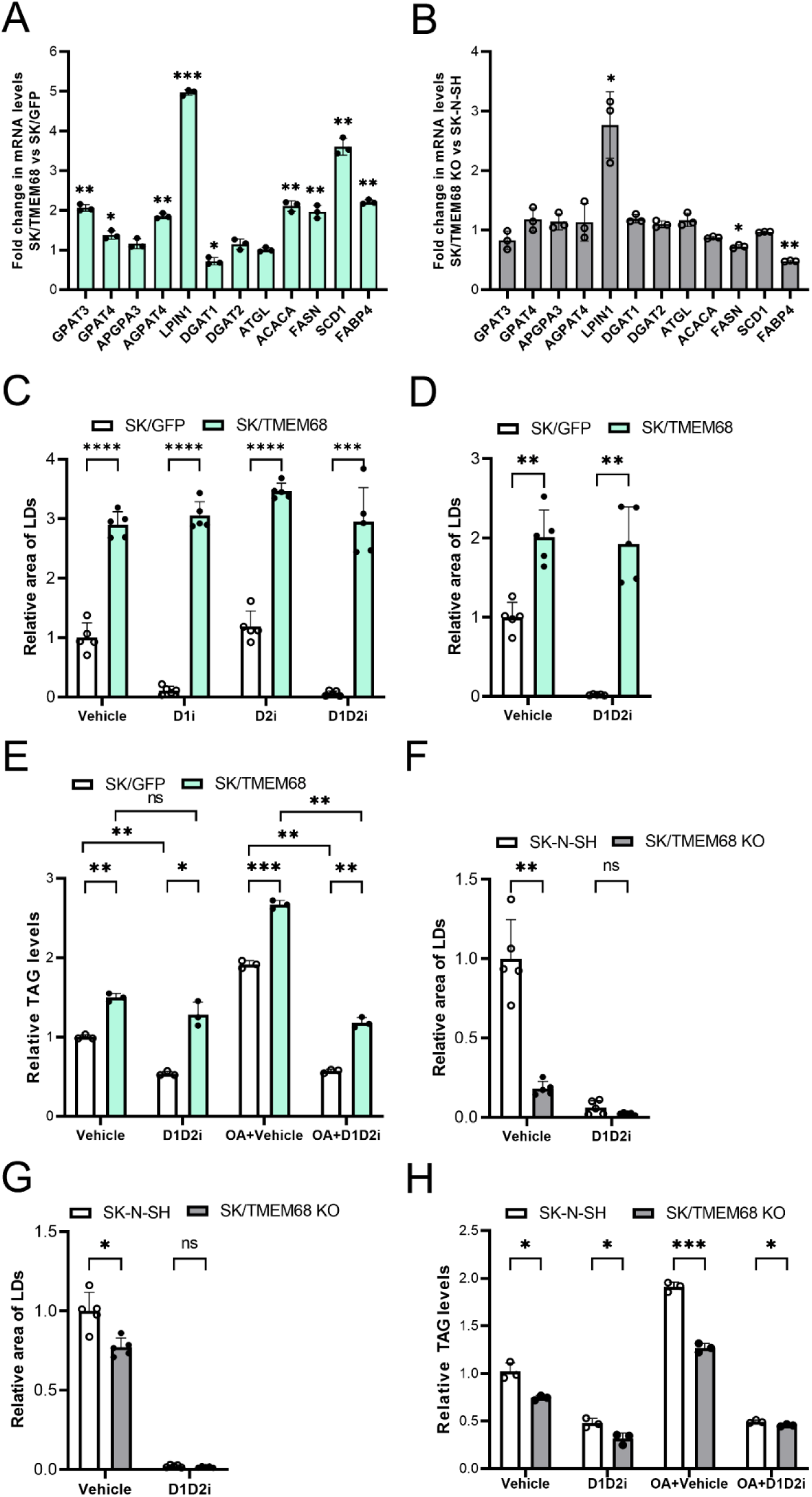
TMEM68 controls TAG independently of DGAT1 and DGAT2. (**A, B**) mRNA levels of genes related to TAG metabolism and fatty acid synthesis in (**A**) SK/TMEM68 and SK/GFP cells; (**B**) SK/TMEM68 KO and SK-N-SH cells. (**C, D**) LD area of (**C**) SK/TMEM68 and SK/GFP cells cultured in the presence of vehicle, DGAT1 inhibitor (D1i), DGAT2 inhibitor (D2i) or D1D2i and in (**C**) the absence or (**D**) presence of OA. (**E**) TAG levels of SK/TMEM68 and SK/GFP cells cultured in the presence of vehicle, D1D2i, and OA as indicated. **(F, G**) LD area in SK/TMEM68 KO and SK-N-SH cells cultured in the presence of vehicle or D1D2i and in (**F**) the absence or (**G**) presence of OA. (**H**) TAG levels of SK/TMEM68 KO and SK-N-SH cells cultured in the presence of vehicle, D1D2i, and OA as indicated. Data are presented as means ± SD. Statistically significant differences are marked with asterisks indicating *P*-values. * *P* < 0.05, ** *P* < 0.01, *** *P* < 0.001, **** *P* < 0.0001, n=3 or *n* = 5. ns, not significant.

Since SK-N-SH express DGAT1 and DGAT2 we wondered, whether TMEM68-mediated TAG formation is independent of DGAT1/2 activity or not. To this end, we blocked DGAT activity pharmacologically using established small molecule inhibitors. The DGAT1 inhibitor (D1i) T863 but not the DGAT2 inhibitor (D2i) PF06424439 methanesulfonate, significantly decreased LDs area by more than 90% in SK/GFP cells as compared to vehicle-treated controls (***Figure 3C***, ***Figure 3-figure supplement 1***), indicating that DGAT1 is the major enzyme for TAG synthesis in SK-N-SH cells. When cells were challenged with both DGAT1 and DGAT2 inhibitors (D1D2i), almost no LDs were observed irrespective of the presence of exogenous OA (***Figure 3C-D***, **Figure 3-figure supplement 1**). In contrast, LD area in SK/TMEM68 cells was largely insensitive to DGAT1/2 inhibition (***Figure 3C-D***, ***Figure 3-figure supplement 1***). In line with our image analyses, exposure to D1D2i significantly decreased TAG levels in basal SK/GFP cells and completely inhibited OA-induced TAG formation (***Figure 3E***). Moreover, D1D2i blocked OA-stimulated TAG synthesis also in SK/TMEM68 cells. Nevertheless, SK/TMEM68 cells exhibited considerable residual TAG levels after exposure to D1D2i, consistent with the abundance of LDs (***Figure 3E*)**. This suggests that TMEM68 is capable of promoting TAG formation independently of DGAT1/2 activities although it does not efficiently metabolize exogenously provided FAs.

We next repeated these experiments with SK/TMEM68 KO cells and found that TMEM68-deficiency was associated with decreased LD area, which was further reduced to undetectable levels by D1D2i treatment (***Figure 3F***, ***Figure 3-figure supplement 2***). Similar results were observed when cells were challenged with exogenous OA, consistent with the notion that incorporation of exogenous FA strictly requires DGATs (***Figure 3G***). Notably, LD area of SK/TMEM68 KO and control cells was indistinguishable after D1D2i treatment (***Figure 3F***). This suggests that endogenous DGAT1/2 essentially define cellular capacity to form LDs with little contribution of TMEM68. Nevertheless, biochemical TAG measurements revealed that SK/TMEM68 KO cells consistently exhibited less cellular TAG as compared to controls even after D1D2i treatment (***Figure 3H***). Thus, while DGAT1 acts as dominant enzyme in SK-N-SH TAG and LD formation our assays disclose a discrete yet consistent basal contribution of endogenous TMEM68 to cellular TAG formation.

### TMEM68 expression alters cellular GPL composition

Our previous study revealed that TMEM68 expression increases MGAT and DGAT activities *in vitro* (*Wang et al., 2023*). However, a recent study concluded that TMEM68 possibly uses GPLs rather than acyl-CoA as acyl donor to convert DAG to TAG (*McLelland et al., 2023*). This prompted us to analyze how TMEM68 affects GPLs in our cellular models. The total levels of PC and PE, which are the most abundant cellular GPL classes, were not significantly affected by TMEM68 overexpression (***Figure 4A***). However, the abundance of ether-linked alkyl-(1-O-alkyl-2-acyl-, i.e., “O”) PC (PC-O) and PE (PE-O) decreased by about 50% and 35%, respectively, in SK/TMEM68 cells compared to controls (***Figure 4A***). Similar changes were observed in subgroups of PC, PE, PC-O, and PE-O, in which FAs at the *sn*-1 and *sn*-2 positions could be fully defined with our analyses (PC-FA, PE-FA, PC-O-FA, and PE-O-FA, ***Figure 4B***). In addition, several anionic GPL classes were altered in SK/TMEM68 cells including decreased levels of bis(monoacylglycero)phosphate (BMP) (about 30%), PG and PI (both about 20%), and increased levels of PA (about 2.7-fold) (***Figure 4A***). Moreover, lysophosphatidylcholine (LPC) but not lysophosphatidylethanolamine (LPE) was decreased by ∼60% upon TMEM68 overexpression (**Figure 4C**). The analysis of individual GPL species revealed that TMEM68 overexpression broadly reduced PC-O and PE-O species containing SFAs, MUFAs, and PUFAs at the *sn*-2 position with little bias towards a particular species (***Figure 4D*, *4E***). Similarly, both MUFAs and PUFAs at the sn-1 position of PC-O and PE-O were reduced in SK/TMEM68 compared to SK/GFP cells (***Figure 4-figure supplement 1***). Despite unaltered overall cellular PC and PE content the molecular composition of those lipid classes differed between SK/TMEM68 and SK/GFP cells. Specifically, PC species with 2-5 double bonds at the *sn*-2 position were reduced while PC species with SFAs and MUFAs at the *sn*-2 position were increased upon TMEM68 overexpression arguing for decrease in overall PC saturation (***Figure 4F***). Similarly, PE species containing FAs with 2 double bonds at the sn-2 position were increased at the expense of >2 double bonds (***Figure 4G***). Changes at the sn-1 position were less consistent but involved a decrease in SFAs in PC and PE (***Figure 4-figure supplement 1***). A relative reduction in PUFAs at the sn-2 position was also observed in other GPLs including PS, PG, PI and BMP (***Figure 4-figure supplement 2***). Together, these analyses suggest that TMEM68 expression depletes ether-linked PC-O and PE-O and strongly alters GPL composition towards higher saturation.

**Figure 4.**
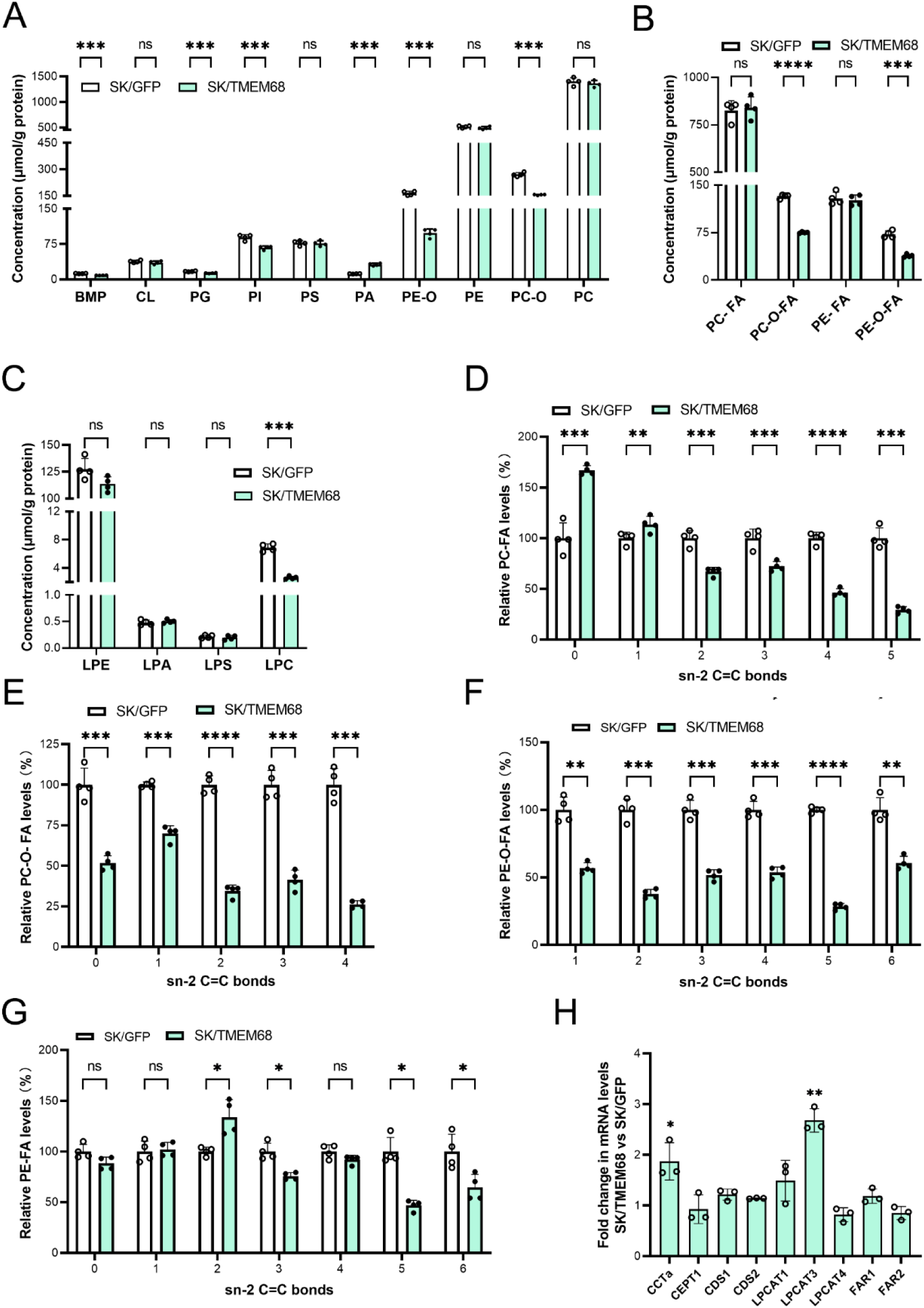
Impact of TMEM68 overexpression on GLPs. (**A**) Total GPL levels in SK/TMEM68 and SK/GFP cells. (**B**) PC-FA, PC-O-FA, PE-FA, and PE-O-FA levels in SK/TMEM68 and SK/GFP cells. (**C**) Levels of total LysoGPLs in SK/TMEM68 and SK/GFP cells. (**D-G**) Relative prevalence of double bonds at the sn-2 position of (**D**) PC-O (**E**), PE-O (**F**), PC, and (**G**) PE in SK/TMEM68 and SK/GFP cells. (**H**) mRNA expression of genes involved in GLP metabolism in SK/TMEM68 and SK/GFP cells. Data are presented as means ± SD. Statistically significant differences are marked with asterisks indicating *P*-values. * *P* < 0.05, ** *P* < 0.01, *** *P* < 0.001, **** *P* < 0.0001, *n* = 4 or n=3. ns, not significant.

To investigate, whether alterations in GPLs are associated with changes in the mRNA level of GPL metabolizing enzymes we performed RT-qPCR. CTP:phosphocholine cytidylyltransferase α (CCTα), encoding for the rate-limiting enzyme in PC synthesis, was expressed at higher levels in SK/TMEM68 compared to SK/GFP cells while expression of choline/ethanolamine phosphotransferase 1 (CEPT1) was unchanged (***Figure 4H*)**. Likewise, expression of CDP-DAG synthase genes *CDS1* and *CDS2*, responsible for the synthesis of anionic GPLs, was unchanged by TMEM68 overexpression. Notably, LPC acyltransferase 3 (LPCAT3), which catalyzes incorporation of PUFAs into PC, was induced in SK/TMEM68 cells (***Figure 4H*)** indicating a compensatory cellular response to reduced PUFAs in GPLs. Given the depletion of ether-linked GPLs we finally assessed expression of FA reductase 1 and 2 (FAR1, FAR2), which generate fatty alcohol precursors for ether lipid synthesis. However, both FARs were expressed at similar levels in SK/TMEM68 and SK/GFP cells (***Figure 4H*)**.

To understand the impact of TMEM68-deficiency on GPLs we repeated these analyses using our SK/TMEM68 KO model. Total levels of most GPL classes were unaltered by TMEM68-deficiency the exception being moderate reductions in PC-O and LPC levels (16% and 23%) **(*Figure 5A*, *5C***). In addition, levels of PC-FA, PE-FA as well as their ether forms PC-O-FA and PE-O-FA were not significantly altered **(*Figure 5B*)**. However, despite largely preserved total GPL levels TMEM68-deficiency altered GPL composition in a manner resembling TMEM68 overexpression. Specifically, PC, PC-O, PE, and PE-O pools were more saturated at the sn-2 position in SK/TMEM68 KO cells compared to controls **(*Figure 5D-5G***). PC and PC-O displayed a significant decrease in species with two or more double bonds at the sn-2 position and a tendency for a relative increase in MUFAs and SFAs, respectively **(*Figure 5D***, ***5E***). Similar trends were observed for PE-O and PE, which exhibited relative increases of sn-2 FAs with one and two double bonds, respectively, at the expense of less saturated FAs **(*Figure 5F***, ***5G*)**. Other GLP classes including PS, PG, PI and BMP showed a similar tendency for a relative reduction of PUFAs at the sn-2 position in SK/TMEM68 KO compared to controls (***Figure 5-figure supplement 1***). To assess whether altered GPLs were associated with dysregulation of genes involved in GPLs or ether lipid synthesis we repeated our RT-qPCR analyses. However, we found little difference in mRNA expression of those genes between SK/TMEM68 KO and SK-N-SH cells **(*Figure 5H*)**. Taken together, these analyses revealed that TMEM68-deficiency, besides reduced lipid storage, is associated with increased saturation of specific GPL classes.

**Figure 5.**
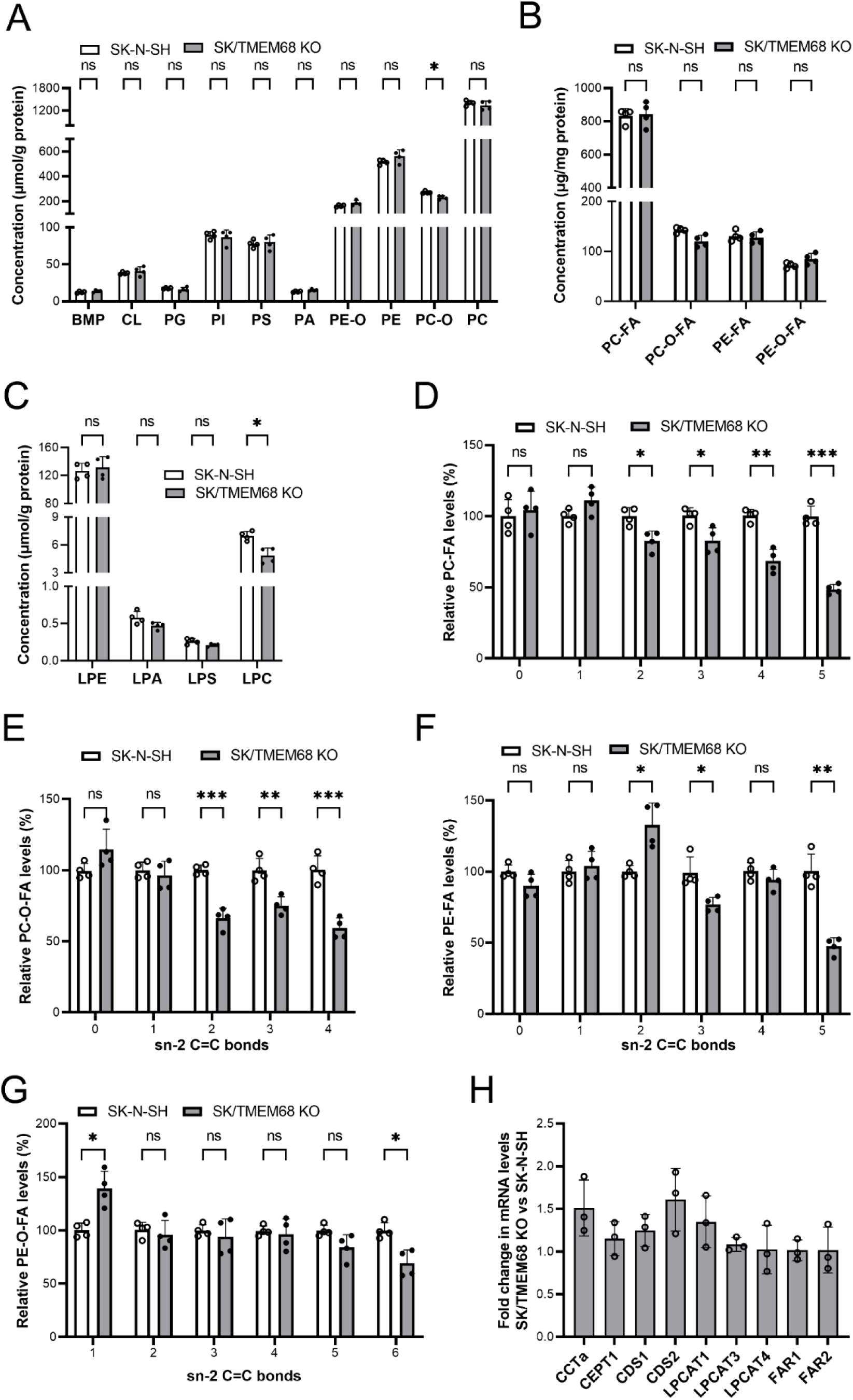
Impact of TMEM68 deficiency on GPLs. (A) Total GPL levels in SK/TMEM68 KO and SK-N-SH cells. (**B**) PC-FA, PC-O-FA, PE-FA, and PE-O-FA levels in SK/TMEM68 KO and SK-N-SH cells. (**C**) Levels of total LysoGPLs in SK/TMEM68 KO and SK-N-SH cells. (**D-G**) Relative prevalence of double bonds at the *sn*-2 position of (**D**) PC-O, (**E**) PE-O, (**F**) PC, and (**G**) PE in SK/TMEM68 KO and SK-N-SH cells. (**H**) mRNA expression of genes involved in GLP metabolism in SK/TMEM68 KO and SK-N-SH cells. Data are presented as means ± SD. Statistically significant differences are marked with asterisks indicating *P*-values. * *P* < 0.05, ** *P* < 0.01, *** *P* < 0.001, n = 4 or n=3. ns, not significant.

### TMEM68 expression affects sphingolipids levels

Although sphingolipids are vital for all eukaryotic cells they are relatively abundant in neuronal cells. We therefore analyzed several sphingolipid classes and found that levels of ceramide trihexoside (Gb3), monosialoganglioside (GM3), ceramide (Cer), hexosylceramide (HexoCer) and lactosylceramide (LacCer) were decreased in SK/TMEM68 compared to SK/GFP cells. In contrast, sphingomyelin, the most abundant sphingolipid, was unaffected by TMEM68-overexpression (***Figure 6A*)**. Notably, SK/TMEM68 KO cells displayed also a reduction in Gb3, HexoCer, LacCer, GM3 and Cer as compared to controls though at lesser extent than SK/TMEM68 cells (***Figure 6B*)**. Taken together, these data show that altering TMEM68 expression disturbs sphingolipid metabolism in neuroblastoma cells.

**Figure 6.**
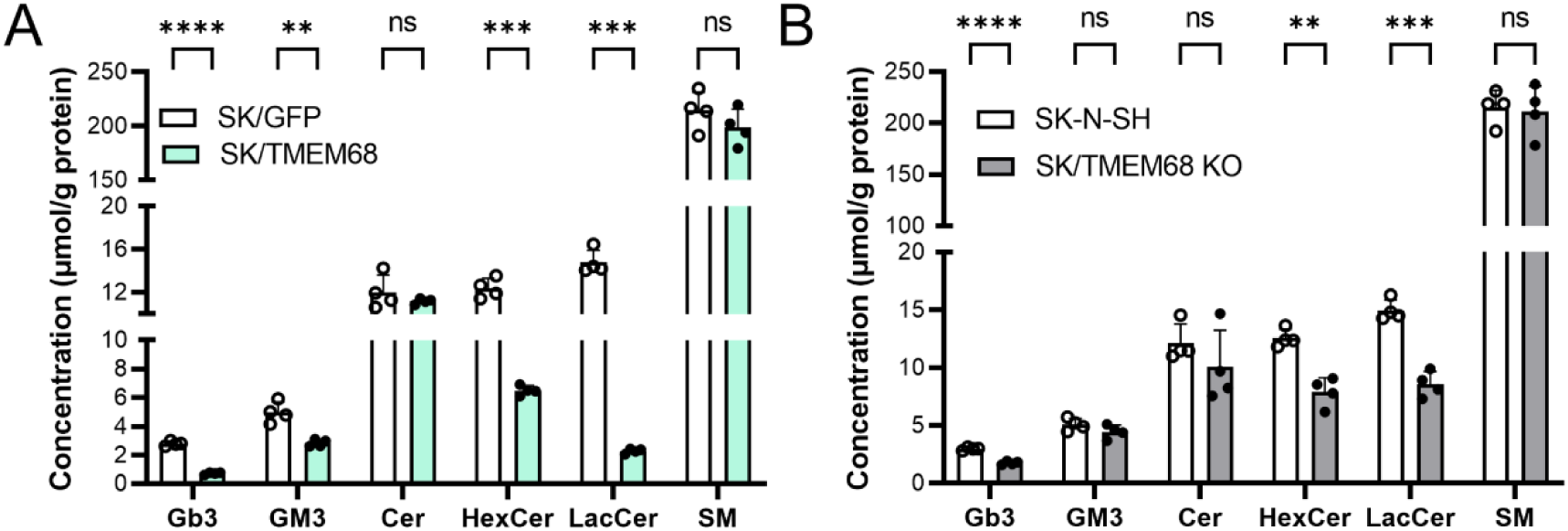
Impact of TMEM68 on sphingolipids. (**A**, **B**) Total sphingolipid levels in (**A**) SK/GFP and SK/TMEM68 cells; (**B**) SK-N-SH and SK/TMEM68 KO cells. Data are presented as means ± SD. Statistically significant differences are marked with asterisks indicating *P*-values. ** *P* < 0.01, *** *P* < 0.001, **** *P* < 0.0001, n = 4. ns, not significant.

## Discussion

The enzymatic synthesis of TAG for storage in intracellular LDs is fundamental to lipid homeostasis, energy metabolism, and stress resistance of eukaryotic cells (*Shi and Cheng, 2009*; *Stone, 2022*). TMEM68 has been previously identified by us and others as a novel ER-anchored TAG synthase (*Wang et al., 2023*; *McLelland et al., 2023*). In the present study we identified additional roles of TMEM68 in cellular GPL and PUFA metabolism.

TMEM68 overexpression robustly increased TAG and LD formation in SK-N-SH neuroblastoma and U251 gliablastoma cells. Pharmacological DGAT1 and DGAT2 inhibition did not compromise the ability of TMEM68 to promote TAG suggesting that TMEM68 operates autonomously and constitutes a novel TAG synthesis pathway in mammalian cells. While pharmacological inhibitors clearly established DGAT1 as major TAG and LD forming enzyme in neuroblastoma cells, a small but discrete portion of basal cellular TAG synthesis was attributable to endogenous TMEM68. Thus, TMEM68 appears to operate in a non-redundant manner with DGAT1 (and DGAT2) in neuroblastoma TAG synthesis and provides a limited yet discrete fraction of basal cellular TAG. Our experiments revealed that TMEM68 differs from DGAT1 in terms of substrate selection. Exogenous FAs were efficiently channeled into TAG by DGAT1 yet were apparently inaccessible to TMEM68. Moreover TMEM68-deficiency caused a biased depletion of highly unsaturated TAG species suggesting that endogenous TMEM68 preferentially (but not exclusively) incorporates PUFAs compared to MUFAs and SFAs. This pattern may reflect differential access of TMEM68 and DGAT1 to precursor pools and could explain why TMEM68 cannot compensate for loss of DGAT1 activity. Interestingly, McLelland *et al*. recently revealed that TMEM68 associates with TMX1, which suppresses its basal activity in several cell lines (*McLelland et al., 2023*). Knockout of TMX1 unleashed TMEM68 activity and increased cellular TAG above basal levels (*McLelland et al., 2023*). Although not formally tested, it is plausible that TMX1 limits TMEM68 in our cell model as well, which would additionally compromise the capacity of TMEM68 to substitute for DGATs.

Based on a weak homology of TMEM68 to DGAT2 we initially hypothesized that TMEM68 possesses MGAT or DGAT activity. In line with this notion, overexpression of TMEM68 in HEK293 cells increased TAG levels along with cellular MGAT and DGAT activities (*Wang et al., 2023*). Nevertheless, these enzyme activities were measured in crude cell homogenates and it remains unclear whether they are a genuine feature of TMEM68 or rather reflect altered expression of endogenous MGAT/DGATs. McLelland et al. recently presented evidence that TMEM68 possibly acts as a PDAT rather than a MGAT/DGAT (*McLelland et al., 2023*). This was based on the observation that ectopic TMEM68 expression while increasing cellular TAG depleted specific GPL classes, in particular ether-linked PC. A formal proof of this proposed activity by means of an in vitro assay however, has not been established. Nevertheless, our present study adds further support to the concept that TMEM68 metabolizes GPLs. Concomitant with increased TAG levels TMEM68 overexpression depleted PC-O and PE-O and - to a lesser extent - several anionic GPLs. Moreover, across multiple GPL classes ectopic TMEM68 expression decreased polyunsaturated at the expense of less unsaturated species. While these data clearly argue for GPLs – in particular ether GPLs – as substrates for TMEM68 the complexity of lipidomic alterations in our system makes it difficult to unequivocally assign clear product-substrate relationships and deduce an underlying enzyme activity. *In vitro* assays with purified or semi-purified preparations are thus required to determine enzymatic activities of TMEM68 and discriminate direct actions of the enzyme from indirect cellular adaptations.

Consistent with a constitutive role of the enzyme in GPL metabolism TMEM68-deficiency elicited several changes in the cellular membrane lipid pattern. While the overall cellular levels of major GPLs remained mostly unaffected by TMEM68-deficiency, their molecular composition was significantly altered. Specifically, GPL species containing PUFAs at the sn-2 position exhibited a relative reduction at the expense of less unsaturated FAs. Thus, in addition to TAG synthesis endogenous TMEM68 appears to have a function in basal cellular GPL remodeling and PUFA homeostasis. De novo synthesized GPLs are typically first esterified to MUFAs and SFAs and get enriched with PUFAs in a hydrolysis/acylation series termed Land’s cycle (*Shindou and Shimizu, 2009*; *Hishikawa et al., 2014)*. To this end, a phospholipase A_2_ hydrolyzes the sn-2 bond of the GPL to generate a lysoGPL, which is subsequently re-acylated with PUFAs by an acyltransferase. PDAT activity also generates lysoGPL besides TAG and can therefore promote cellular GPL catabolism and remodeling (*Yoon et al., 2012; Barbosa et al., 2019*). Given the presumed PDAT function of TMEM68 it is conceivable that its loss would diminish PUFA incorporation in neuroblastoma cells by slowing GPL turnover.

Curiously, both loss and gain of TMEM68 function elicited somewhat similar alterations in membrane lipid composition including depletion of PUFA from GPLs and decreases in specific SPLs. It should be noted, however, that TMEM68 mRNA expression was increased 100-fold over endogenous levels in our overexpression model. Given that TMEM68 acts as a GPL-metabolizing enzyme increasing its levels above a certain threshold may enforce compensatory cellular responses to mitigate the damaging effect of its activity. In line with this notion the mRNA levels of several enzymes of the FA synthesis and G3P pathways were induced upon TMEM68 overexpression and the G3P pathway intermediates PA and DAG were increased. We also observed a disproportionate accumulation of saturated and monounsaturated TAG species in TMEM68-expressing cells likely reflecting an enhanced supply from de novo FA synthesis. Hence, while alterations in GPL composition in TMEM68-deficient cells likely reflect a physiological function of the enzyme the same alterations in TMEM68-overexpressing cells may be a conglomeration of compensatory cellular adaptions to unleashed TMEM68 activity.

In summary, our study provides insights into human TMEM68 functioning as a novel TAG synthase with a substrate spectrum distinct from canonical DGATs and disclose a role of TMEM68 in linking storage lipid metabolism to GPL homeostasis.

## Material and Methods

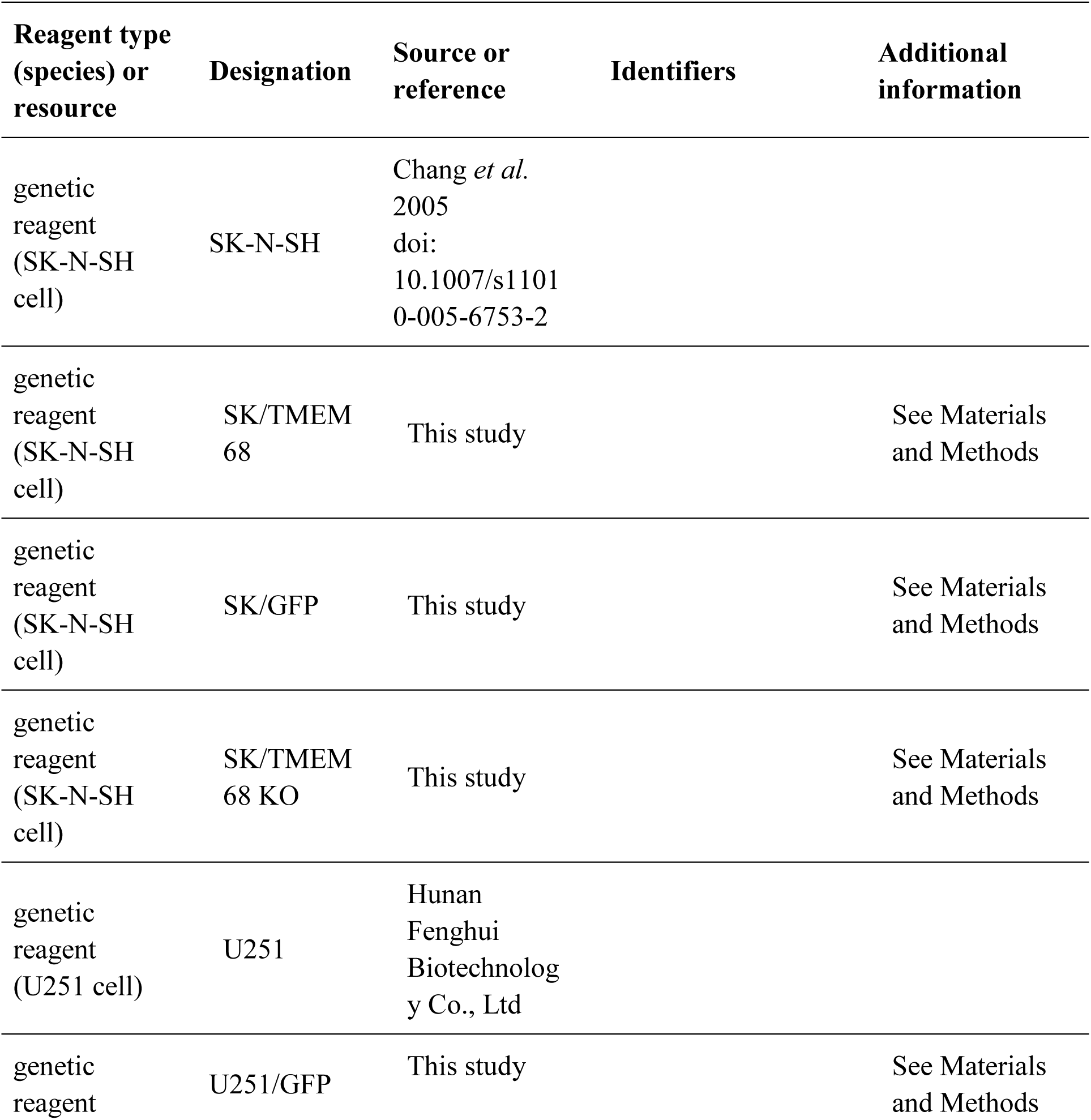
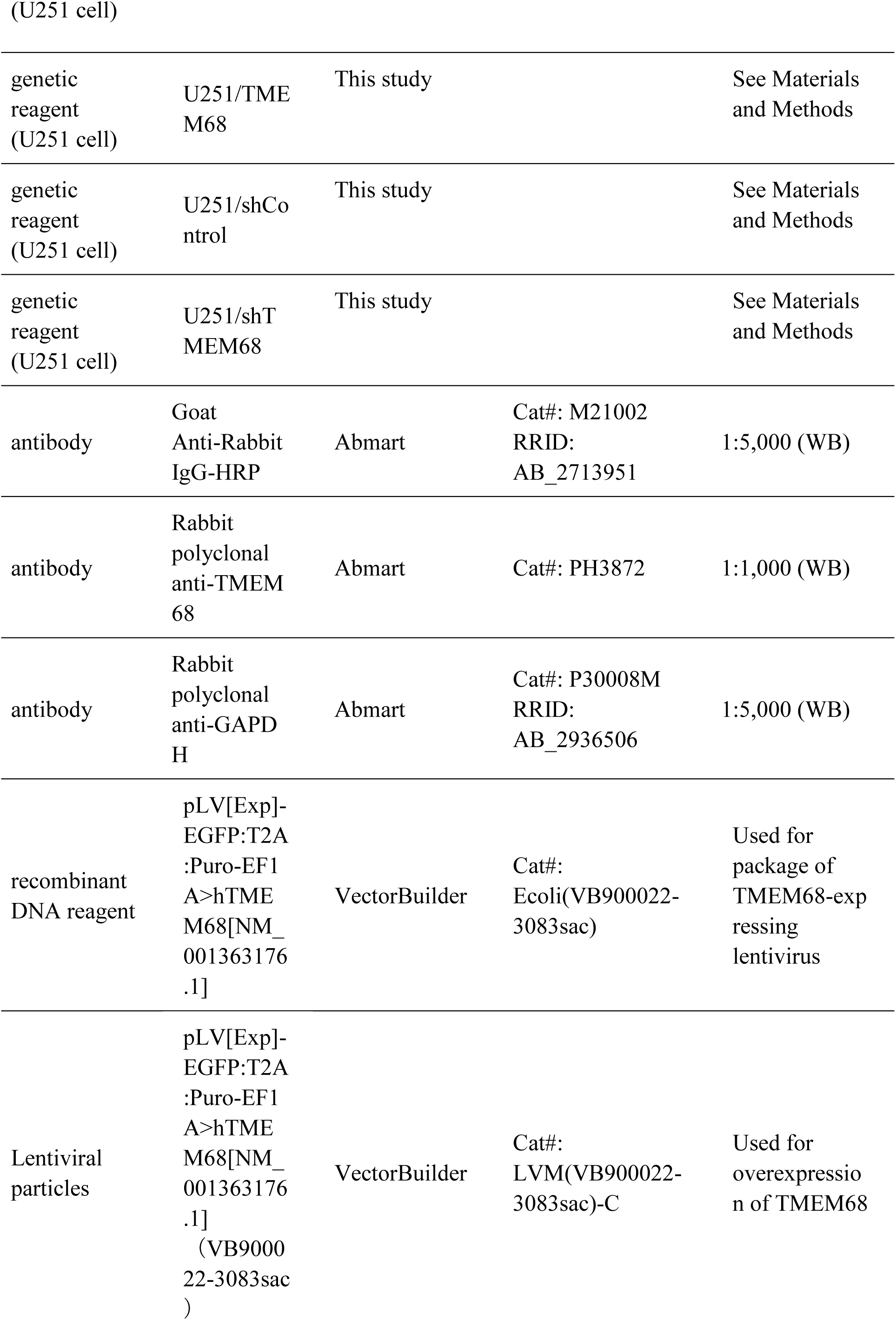
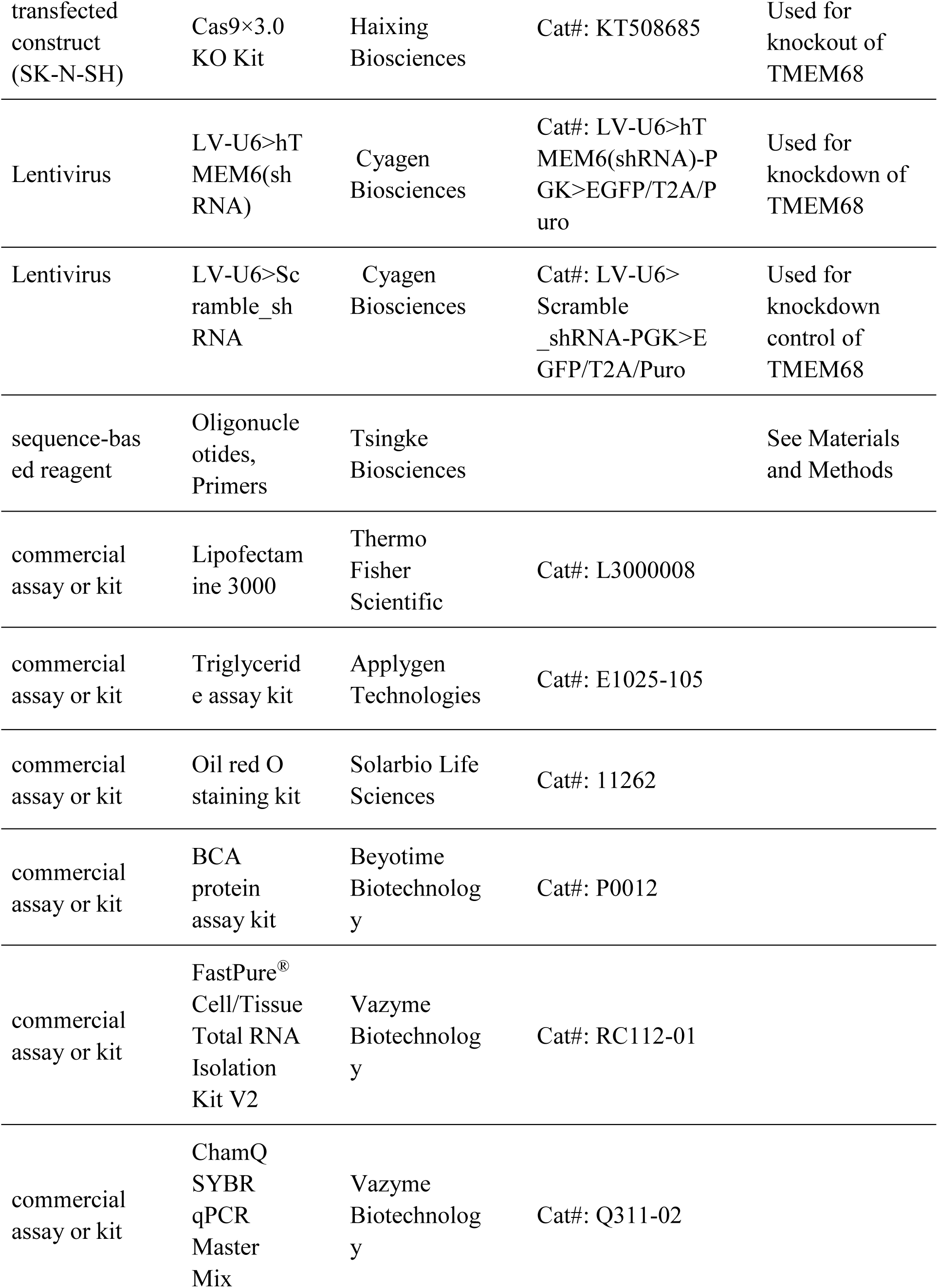
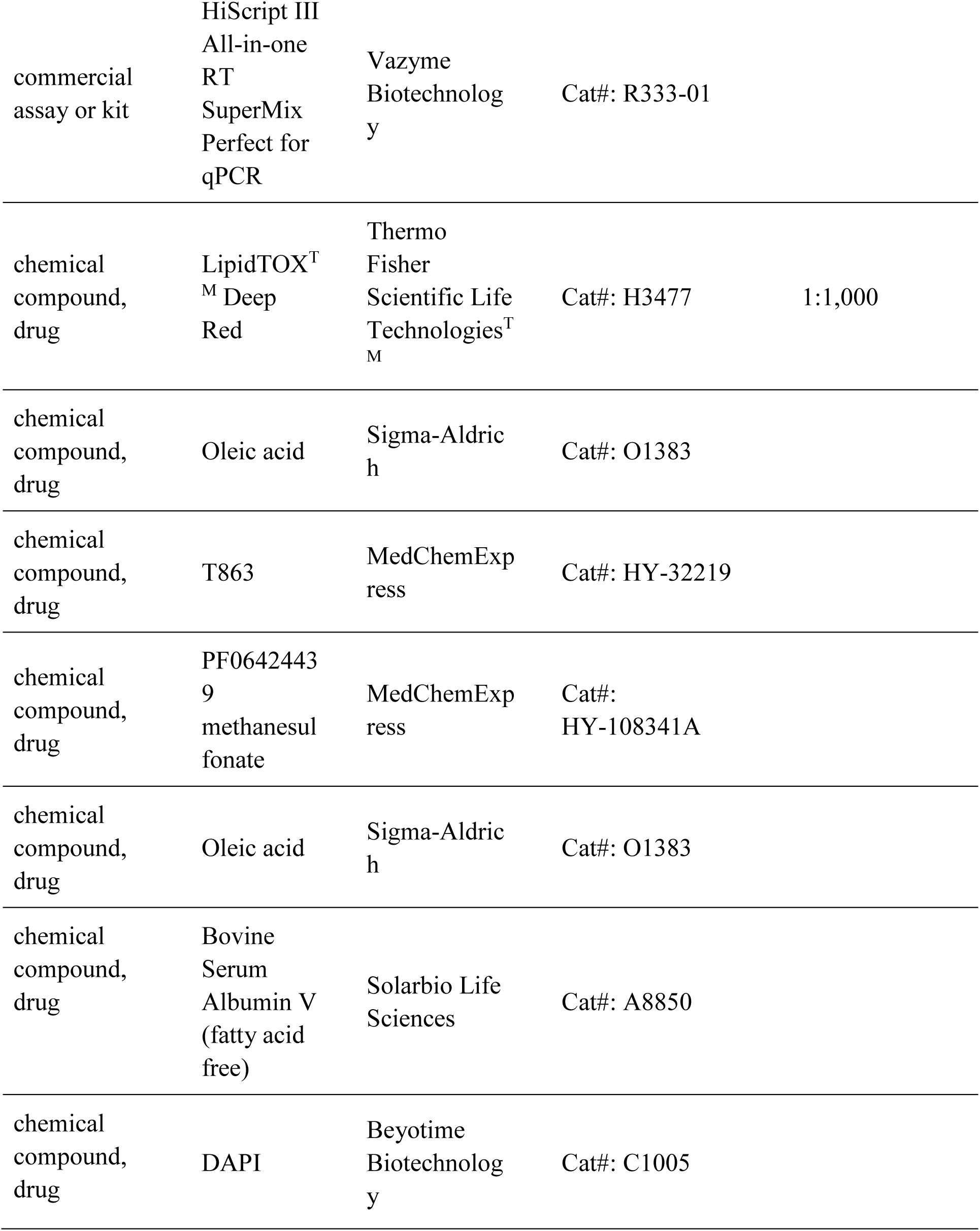
Key Resources Table

Key Resources Table
| Reagent type<br>(species) or<br>resource | Designation | Source or<br>reference | Identifiers | Additional<br>information |
| --- | --- | --- | --- | --- |
| genetic<br>reagent<br>(SK-N-SH<br>cell) | SK-N-SH | Chang *et al.*<br>2005<br>doi:<br>10.1007/s1101<br>0-005-6753-2 |  |  |
| genetic<br>reagent<br>(SK-N-SH<br>cell) | SK/TMEM<br>68 | This study |  | See Materials<br>and Methods |
| genetic<br>reagent<br>(SK-N-SH<br>cell) | SK/GFP | This study |  | See Materials<br>and Methods |
| genetic<br>reagent<br>(SK-N-SH<br>cell) | SK/TMEM<br>68 KO | This study |  | See Materials<br>and Methods |
| genetic<br>reagent<br>(U251 cell) | U251 | Hunan<br>Fenghui<br>Biotechnolog<br>y Co., Ltd |  |  |
| genetic<br>reagent | U251/GFP | This study |  | See Materials<br>and Methods |
| (U251 cell) |  |  |  |  |
| genetic reagent (U251 cell) | U251/TME M68 | This study |  | See Materials and Methods |
| genetic reagent (U251 cell) | U251/shControl | This study |  | See Materials and Methods |
| genetic reagent (U251 cell) | U251/shT MEM68 | This study |  | See Materials and Methods |
| antibody | Goat Anti-Rabbit IgG-HRP | Abmart | Cat#: M21002<br>RRID: AB_2713951 | 1:5,000 (WB) |
| antibody | Rabbit polyclonal anti-TMEM68 | Abmart | Cat#: PH3872 | 1:1,000 (WB) |
| antibody | Rabbit polyclonal anti-GAPDH | Abmart | Cat#: P30008M<br>RRID: AB_2936506 | 1:5,000 (WB) |
| recombinant DNA reagent | pLV[Exp]-EGFP:T2A :Puro-EF1 A>hTME M68[NM_001363176.1] | VectorBuilder | Cat#: Ecoli(VB900022-3083sac) | Used for package of TMEM68-expressing lentivirus |
| Lentiviral particles | pLV[Exp]-EGFP:T2A :Puro-EF1 A>hTME M68[NM_001363176.1] (VB900022-3083sac) | VectorBuilder | Cat#: LVM(VB900022-3083sac)-C | Used for overexpression of TMEM68 |
| transfected construct (SK-N-SH) | Cas9×3.0 KO Kit | Haixing Biosciences | Cat#: KT508685 | Used for knockout of TMEM68 |
| Lentivirus | LV-U6>hT MEM6(sh RNA) | Cyagen Biosciences | Cat#: LV-U6>hT MEM6(shRNA)-PGK>EGFP/T2A/Puro | Used for knockdown of TMEM68 |
| Lentivirus | LV-U6>Scramble_sh RNA | Cyagen Biosciences | Cat#: LV-U6>Scramble_shRNA-PGK>EGFP/T2A/Puro | Used for knockdown control of TMEM68 |
| sequence-based reagent | Oligonucleotides, Primers | Tsingke Biosciences |  | See Materials and Methods |
| commercial assay or kit | Lipofectamine 3000 | Thermo Fisher Scientific | Cat#: L3000008 |  |
| commercial assay or kit | Triglyceride assay kit | Applygen Technologies | Cat#: E1025-105 |  |
| commercial assay or kit | Oil red O staining kit | Solarbio Life Sciences | Cat#: 11262 |  |
| commercial assay or kit | BCA protein assay kit | Beyotime Biotechnology | Cat#: P0012 |  |
| commercial assay or kit | FastPure<sup>®</sup> Cell/Tissue Total RNA Isolation Kit V2 | Vazyme Biotechnology | Cat#: RC112-01 |  |
| commercial assay or kit | ChamQ SYBR qPCR Master Mix | Vazyme Biotechnology | Cat#: Q311-02 |  |
| commercial assay or kit | HiScript III All-in-one RT SuperMix Perfect for qPCR | Vazyme Biotechnology | Cat#: R333-01 |  |
| chemical compound, drug | LipidTOX<sup>T</sup><sub>M</sub> Deep Red | Thermo Fisher Scientific Life Technologies<sup>T</sup><sub>M</sub> | Cat#: H3477 | 1:1,000 |
| chemical compound, drug | Oleic acid | Sigma-Aldrich | Cat#: O1383 |  |
| chemical compound, drug | T863 | MedChemExpress | Cat#: HY-32219 |  |
| chemical compound, drug | PF06424439 methanesulfonate | MedChemExpress | Cat#: HY-108341A |  |
| chemical compound, drug | Oleic acid | Sigma-Aldrich | Cat#: O1383 |  |
| chemical compound, drug | Bovine Serum Albumin V (fatty acid free) | Solarbio Life Sciences | Cat#: A8850 |  |
| chemical compound, drug | DAPI | Beyotime Biotechnology | Cat#: C1005 |  |

### Cell culture and treatment

SK-N-SH and U251 cells were cultured in DMEM supplemented with 10% fetal bovine serum (FBS), 100 units/mL penicillin and 100 µg/mL streptomycin in a 37 *◦*C incubator with 5% CO_2_. To induce TAG biosynthesis, cells were incubated in medium containing 0.2 mM OA supplemented with fat-free BSA for 12 h.

### Generation of lentivirus and stable infection of SK-N-SH and U251 cells

Lentiviral particles harboring vectors for the expression of GFP and human TMEM68 proteins respectively were generated by VectorBuilder (Guangzhou, China). Lentiviral constructs encoding for scramble shRNA or shRNAs targeting human TMEM68 (shTMEM68-1 TTCCTGGATAATGCGTTAAATCTCGAGATTTAACGCATTATCCAGGAA, shTMEM68-2 GAGTTCGAGAAGCCCTAATTACTCGAGTAATTAGGGCTTCTCGAACTC, shTMEM68-3 CCAGCCTGAATGATGAATTTACTCGAGTAAATTCATCATTCAGGCTGG) and lentiviral particles were obtained from Cyagen Biosciences (Suzhou, China). SK-N-SH and U251 cells were seeded into 6-well plates at a density of 300,000 cells/well. The next day cells were incubated with lentivirus-containing supernatants in the presence of 8 μg/mL Polybrene for 24 h. To ensure stable expression, cells were maintained for 7 days in medium containing 2 μg/mL puromycin. Clones were grown to 80-90% confluency and then passaged until enough cells could be harvested for overexpression or knockdown validation by RT‒ qPCR and Western blotting.

### Generation and validation of TMEM68 knockout cells using CRISPR-Cas9

CRISPR technology was utilized to generate genetic TMEM68 knockout (KO) cells using Cas9×3.0 CRISPR Gene KO Kit (Haixing Biosciences, Suzhou, China). Two gRNAs were designed to target the sequence of human TMEM68 gene (Gene ID 137695), gRNA1: TGATTCACATACTCGAAGAA; gRNA2: CACATACTCGAAGAATGGTT. Human neuroblastoma SK-N-SH cells (4×10^5^/well) were seeded in 12-well plates. The next day, each gRNA plasmid (0.5 µg) and donor DNA plasmid (0.5 µg) were combined with 2.5 µL of Lipofectamine 3000 and 2 µL of P3000™ reagent in 200 µL of Opti-MEM, which were incubated at room temperature for 15 min. SK-N-SH culture medium was replaced with the appropriate plasmid-lipofectamine solution, and cells were incubated at 37°C for 48 h. The medium was then replaced with DMEM plus 10% FBS and 2 µg/mL puromycin to generate stable cell clones. The isolated cell clones were diluted to single-cell cultures in 96-well plates. Clones were grown to 80-90% confluency and then passaged until enough cells could be harvested for KO validation by RT‒qPCR and Western blotting.

### Western blotting

Total protein extract was prepared in RIPA buffer supplemented with 1 mM phenylmethanesulfonyl fluoride (PMSF) by sonication, and then centrifuged with 15,000× g for 15 min at 4°C to pellet the cell debris. The supernatant protein concentration was determined with BCA protein assay kit (Beyotime Biotechnology, Shanghai, China). The supernatant was mixed with 5 × SDS loading buffer and boiled for 5 min. All the samples were subjected to SDS/PAGE, transferred to nitrocellulose filters and subjected to immunoblotting analysis using an anti-TMEM68 antibody (1:1000 dilution) and an anti-GAPDH (1:5000) antibody as described previously (*Heier et al., 2017*).

### Oil Red O staining and quantification of LD area

Cells were treated with vehicle (0.1% DMSO), 10 µM DGAT1 inhibitor T863, 5 µM DGAT2 inhibitor PF06424439 methanesulfonate or both DGAT inhibitors for 6 h in the presence or absence of 0.1 mM OA complexed to fat-free BSA. After removing the medium and washing with PBS, the cells were fixed with 4% paraformaldehyde in PBS for 10 min at room temperature and then stained using an Oil Red O staining kit (Solarbio, Beijing, China). Cell images were acquired using an Olympus IX73 inverted microscope. The acquired images were processed manually with ImageJ (Fiji) software. The LD area (2D) was quantitatively determined using the ImageJ (Fiji) Auto Local Threshold tool (Bernsen method).

### Measurement of TAG levels

After removing the medium, the cells were washed with PBS twice, harvested, and then dissolved in lysis buffer (100 µL/1×10^6^ cells). Then, the cell lysate was briefly centrifuged, and the protein concentration of the supernatant was assayed with a BCA protein assay kit (Beyotime Biotechnology, Shanghai, China). The supernatant was further heated at 70°C for 10 min and centrifuged for 5 min at 2000 rpm. The TAG levels in the supernatant were determined with a triglyceride assay kit (Applygen, Beijing, China).

### RNA extraction and RT‒qPCR

Total RNA was extracted using FastPure^®^ Cell/Tissue Total RNA Isolation Kit V2 (Vazyme Biotechnology, Nanjing, China) and then transcribed to cDNA with HiScript III All-in-one RT SuperMix Perfect for qPCR Kit (Vazyme Biotechnology, Nanjing, China). Real-time quantitative PCR (qPCR) was performed with ChamQ SYBR qPCR Master Mix (Vazyme Biotechnology, Nanjing, China) on a Bio-Rad IQ5 qPCR System. The primers used were shown in the supplement Table 1. Relative mRNA levels were quantified according to the ΔΔCt method using *β-actin* as reference gene (*Schmittgen and Livak, 2008*).

### Confocal fluorescence microscopy

For detection of LDs, cells were seeded in 12-well plates mounted onto cover slips. After seeding for 24 h, cells were washed with PBS twice and fixed with 4% paraformaldehyde (PFA) for 30 min at room temperature. LDs were stained with HCS LipidTOX^TM^ Deep Red (1:1000 in PBS) for 30 min. After brief washing with PBS, nuclei were stained with DAPI (0.5 µg/mL) for 15 min. Slides were sealed with antifade mounting medium and stored at 4°C. Fluorescent images were captured with a Olympus SpinSR confocal microscope. HCS LipidTOX^TM^ Deep Red was excited at 633 nm, and emission was detected between 650 and 700 nm. DAPI was excited at 364 nm, and emission was detected between 450 and 490 nm. All the presented experiments were repeated independently at least 3 times.

### Quantitative untargeted lipidomic analysis

Quantitative untargeted lipidomic experiments were performed in human neuroblastoma SK-N-SH cells and TMEM68 knockout cells, GFP-expressing cells and TMEM68-expressing cells. Lipids were extracted from approximately one million cells using a modified version of the Bligh and Dyer’s method as described previously (*Song et al., 2020*). Briefly, cells were homogenized in 750 µL of chloroform:methanol:MilliQ H_2_O (3:6:1) (v/v/v). The homogenate was then incubated at 1500 rpm for 1 h at 4 ℃. At the end of the incubation, 350 µL of deionized water and 250 µL of chloroform were added to induce phase separation. The samples were then centrifuged and the lower organic phase containing lipids was extracted into a clean tube. Lipid extraction was repeated once by adding 450 µL of chloroform to the remaining cells in aqueous phase, and the lipid extracts were pooled into a single tube and dried in the SpeedVac under OH mode. Samples were stored at -80℃ until further analysis. Upper aqueous phase and cell pellet were dried in a SpeedVac under H_2_O mode. Total protein content was determined from the dried pellet using the Pierce^®^ BCA Protein Assay Kit according to the manufacturer’s protocol.

Lipidomic analyses were conducted at LipidALL Technologies (Changzhou, China) using a Jasper HPLC system coupled with Sciex TRIPLE QUAD 4500 MD system as reported previously (*Lam et al., 2021*). Separation of individual lipid classes by normal phase (NP)-HPLC was carried out using a TUP-HB silica column (i.d. 150×2.1 mm, 3 µm) under the following conditions:mobile phase A (chloroform:methanol:ammonium hydroxide, 89.5:10:0.5) and mobile phase B (chloroform:methanol:ammonium hydroxide:water, 55:39:0.5:5.5). MRM transitions were set up for comparative analysis of various lipids. Individual lipid species were quantified by referencing spiked internal standards.

### Quantitative targeted free fatty acid analysis

A total of 1×10^6^ cells were collected and resuspended in 100 μL of ultrapure water extract. Half of the cell suspension was mixed well with 75 μL of methanol solution, 100 μL of methyl tert-butyl ether solution, and 25 μL of 36% phosphoric acid solution through vortexing for 3 min. Then, the cell suspension was frozen in liquid nitrogen for 2 min, thawed on ice for 5 min and vortexed for 3 min, which was repeated twice. The lipid extract was centrifuged at 12000 rpm for 5 min at 4°C. Two hundred microlitres of supernatant was dried on a nitrogen blower, 300 μL of 15% boron trifluoride methanol solution was added, the mixture was vortexed for 3 minutes, and the mixture was incubated in an oven at 60°C for 30 minutes. After cooling to room temperature, 500 μL of n-hexane solution and 200 μL of saturated sodium chloride solution were accurately added and mixed well. One hundred microlitres of the n-hexane layer solution was transferred for further analysis after centrifugation at 12000 rpm at 4°C for 5 minutes. Fatty acids and their metabolites were detected by MetWare (Wuhan, China, http://www.metware.cn/) using a GC-EI-MS system (GC, Agilent 8890; MS, 5977B System). The remaining 50 μL of cell suspension was frozen and thawed 3 times and centrifuged at 12,000 r/min for 10 min, after which the protein concentration of the supernatant was determined with a BCA protein assay kit.

### Statistical analysis

Data were generally expressed as mean ± standard deviation (SD) values. Groups of data were compared by one-way ANOVA analysis using the T-TEST method with two tails and two-sample heteroscedasticity. A difference between means was considered significant at *P* < 0.05.

## Author contributions

Fansi Zeng, Data curation, Formal analysis, Validation, Investigation, Visualization, Methodology; Christoph Heier, Methodology, Visualization, Writing – original draft, Writing – review and editing; Qing Yu, Data curation, Methodology, Investigation; Huimin Peng, Formal analysis, Funding acquisition, Project administration; Feifei Huang, Validation, Visualization; Zheng Zhao, Methodology, Investigation; Pingan Chang, Conceptualization, Data curation, Supervision, Investigation, Funding acquisition, Writing – original draft, Project administration, Writing – review and editing.

## Acknowledgements

We thank Chengrong Ling and Wenxia Zheng for helping with confocal fluorescence microscopy. This work was supported by Chongqing Science and Technology Bureau grant CSTB2022NSCQ-MSX0957 (to PC), and in part by the Science and Technology Research Program of Chongqing Municipal Education Commission (Grant No. KJQN20230065 to HP).

## Declaration of interests

The authors declare no competing interests.

## Figures

**Figure 1-figure supplement 1.**
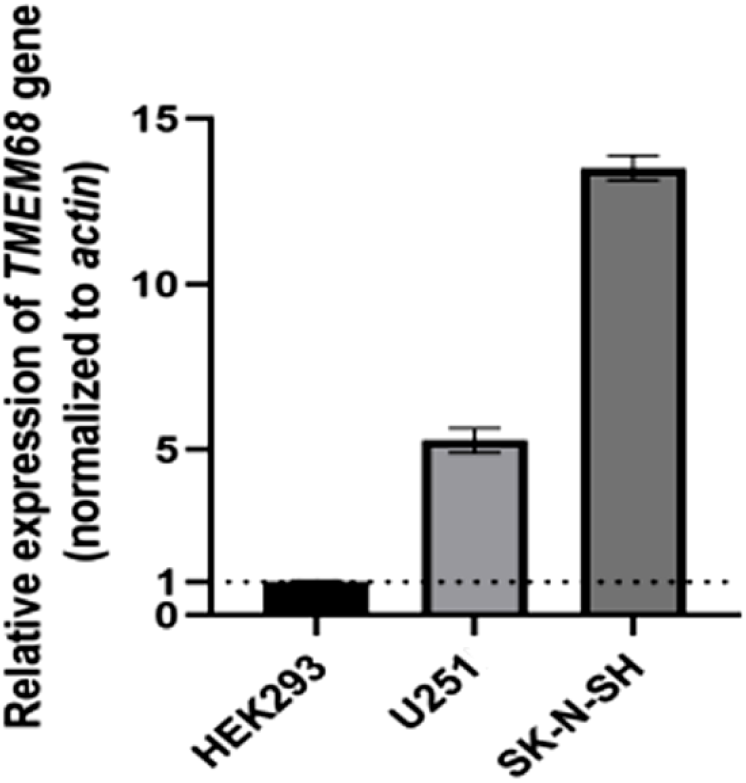
Relative mRNA expression levels of the *TMEM68* gene in three cell lines. Data is presented as means ±SD,*n* = 3.

**Figure 1-figure supplement 2.**
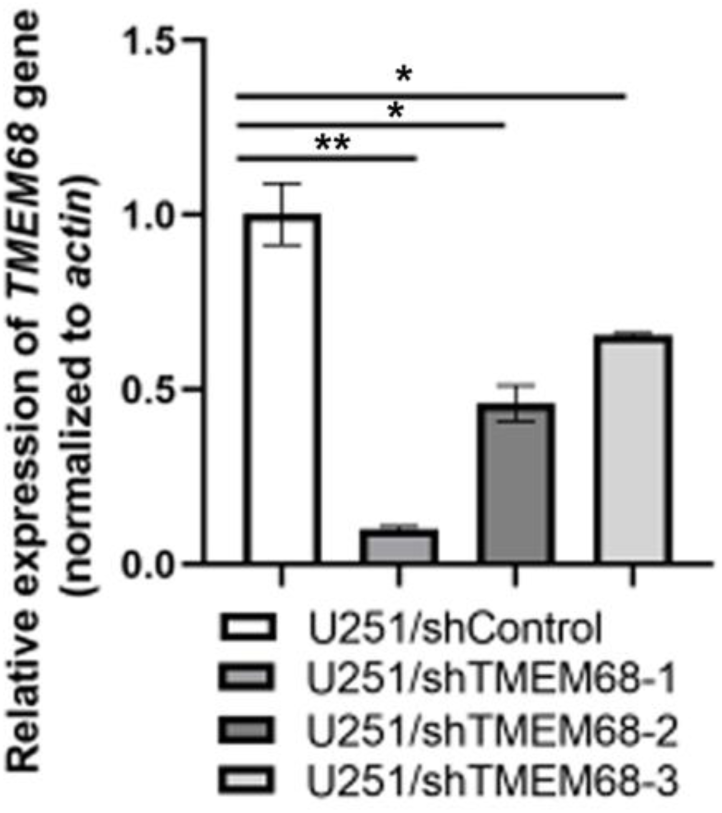
Relative mRNA expression of the *TMEM68* gene in U251 cells expressing one of three different TMEM68 shRNA constructs (U251/shTMEM68-1, -2, -3) or scramble shRNA (U251/shControl), respectively. Data is presented as means ± SD. Asterisk indicates *P-*values: * *P* < 0.05, ** *P* < 0.01. *n* = 3.

**Figure 2-figure supplement 1.**
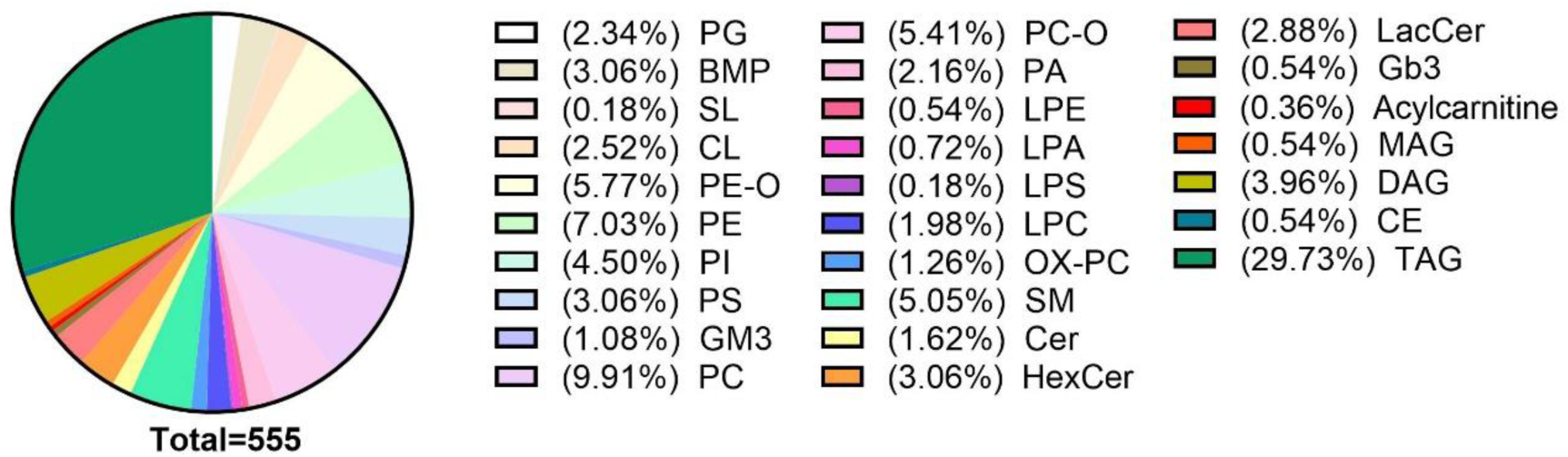
Distribution of 555 detected lipid species detected by LC/MS among different lipid classes.

**Figure 2-figure supplement 2.**
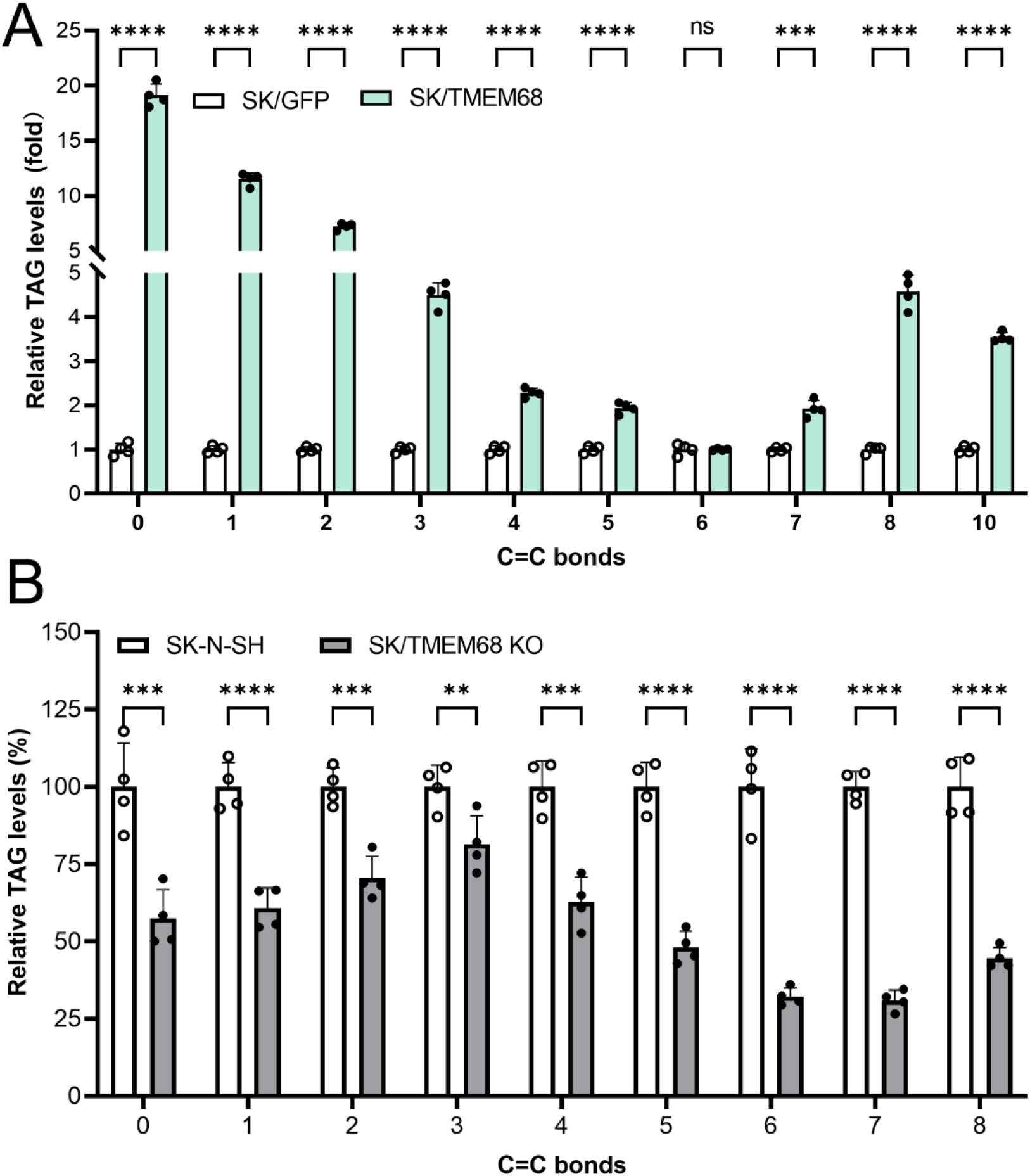
Relative levels of TAG species with different fatty acyl saturation (expressed as total C=C bonds per TAG) (**A**) SK/TMEM68 and SK/GFP control cells; (**B**) SK/TMEM68 KO and SK-N-SH control cells. Data is presented as means ± SD. Statistically significant differences are marked with asterisks indicating *P*-values. ** *P* < 0.01, *** *P* < 0.001, **** *P* < 0.0001. ns, not significant. *n* = 4.

**Figure 2-figure supplement 3.**
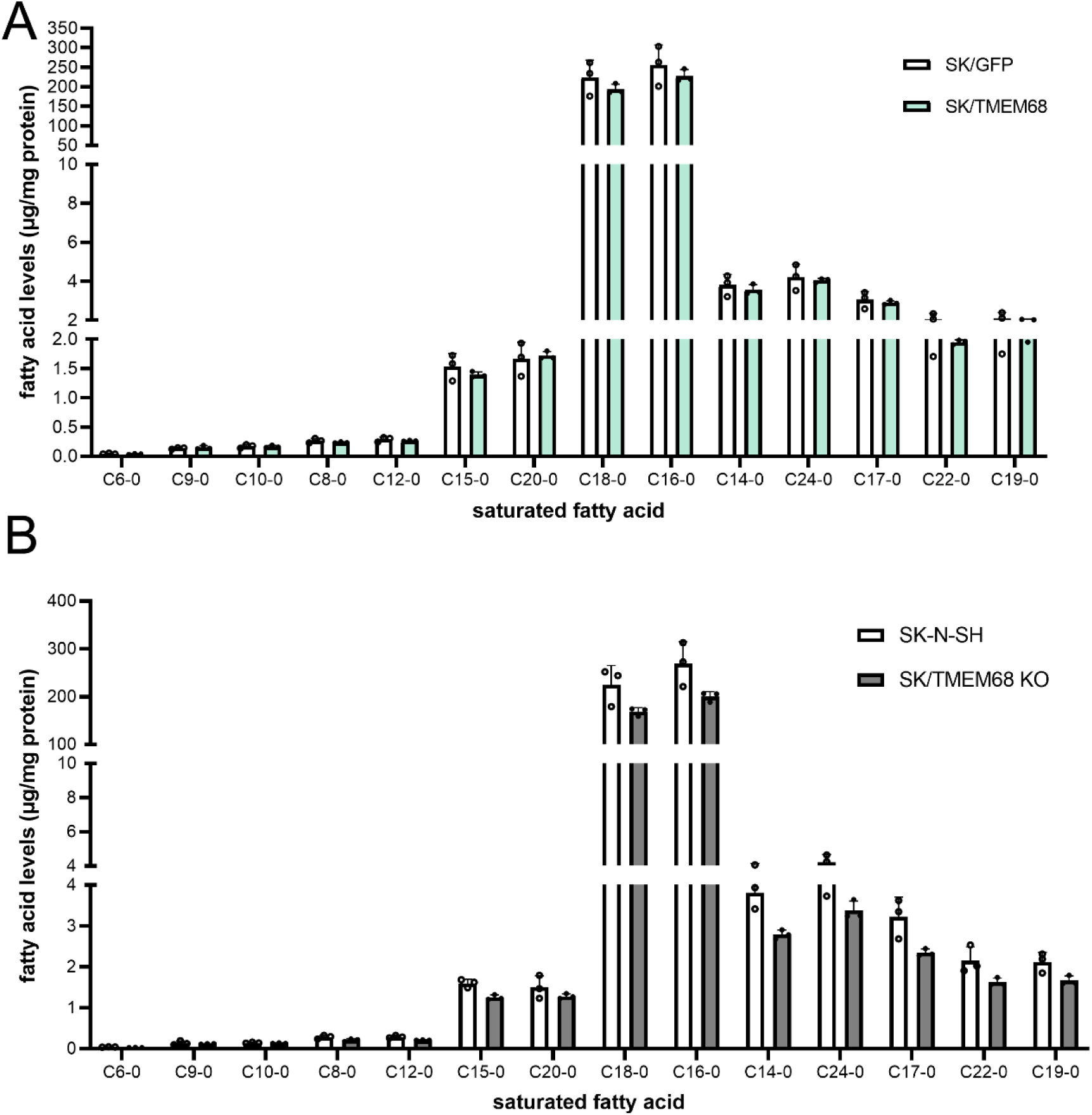
Absolute levels of non-esterified FAs in (**A**) SK/TMEM68 and SK/GFP control cells; (**B**) SK/TMEM68 KO and SK-N-SH control cells. Data is presented as means ± SD. Statistically significant differences are marked with asterisks indicating *P*-values. ** *P* < 0.01, *** *P* < 0.001, **** *P* < 0.0001. ns, not significant. *n* = 4.

**Figure 3-figure supplement 1.**
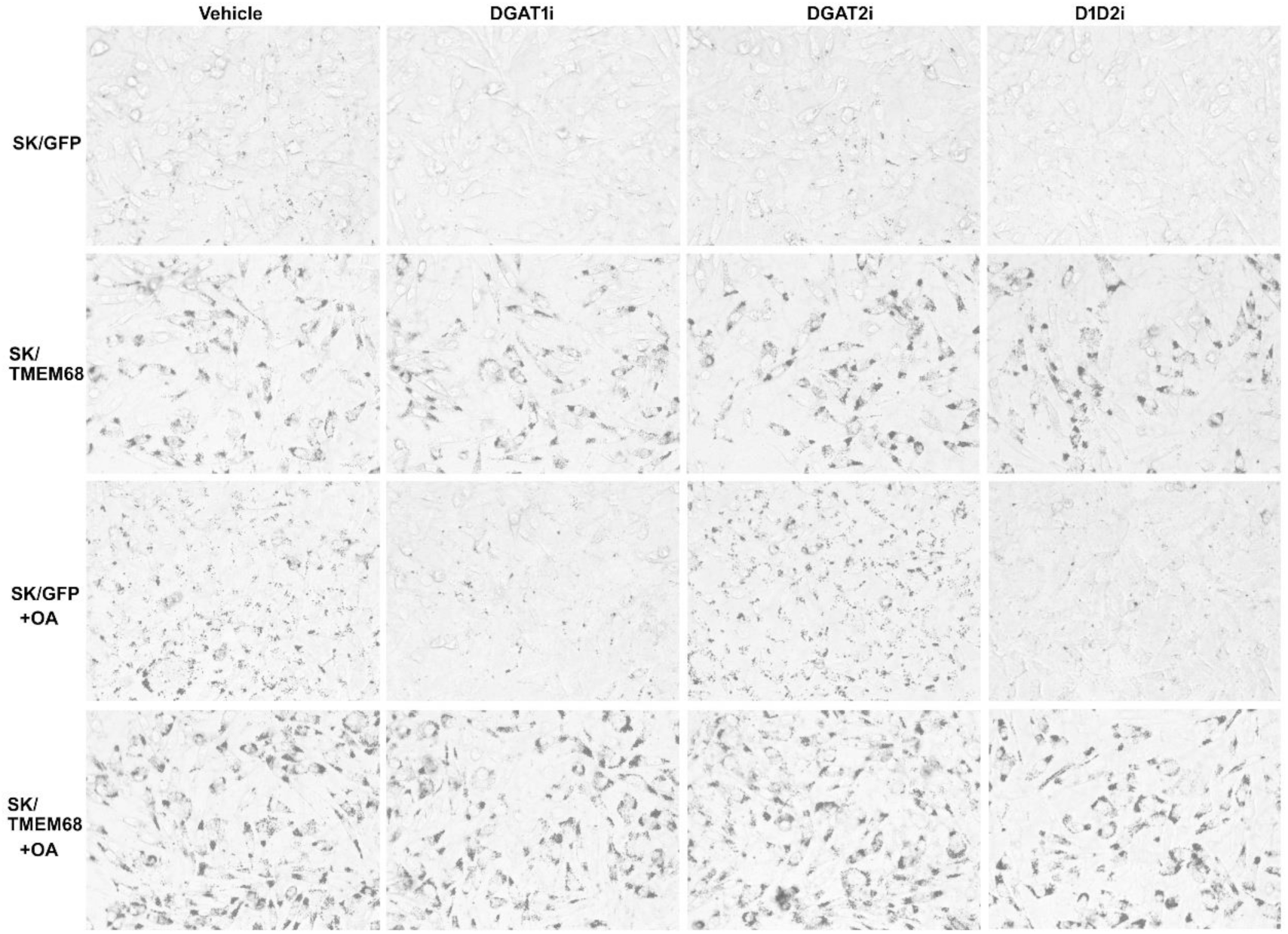
Oil Red O staining of LDs in SK/GFP and SK/TMEM68 cells cultured in in the absence or presence of vehicle, DGAT1 inhibitor (DGAT1i), DGAT2 inhibitor (DGAT2i), a combination of both inhibitors (D1D2i), and OA (+OA) as indicated.

**Figure 3-figure supplement 2.**
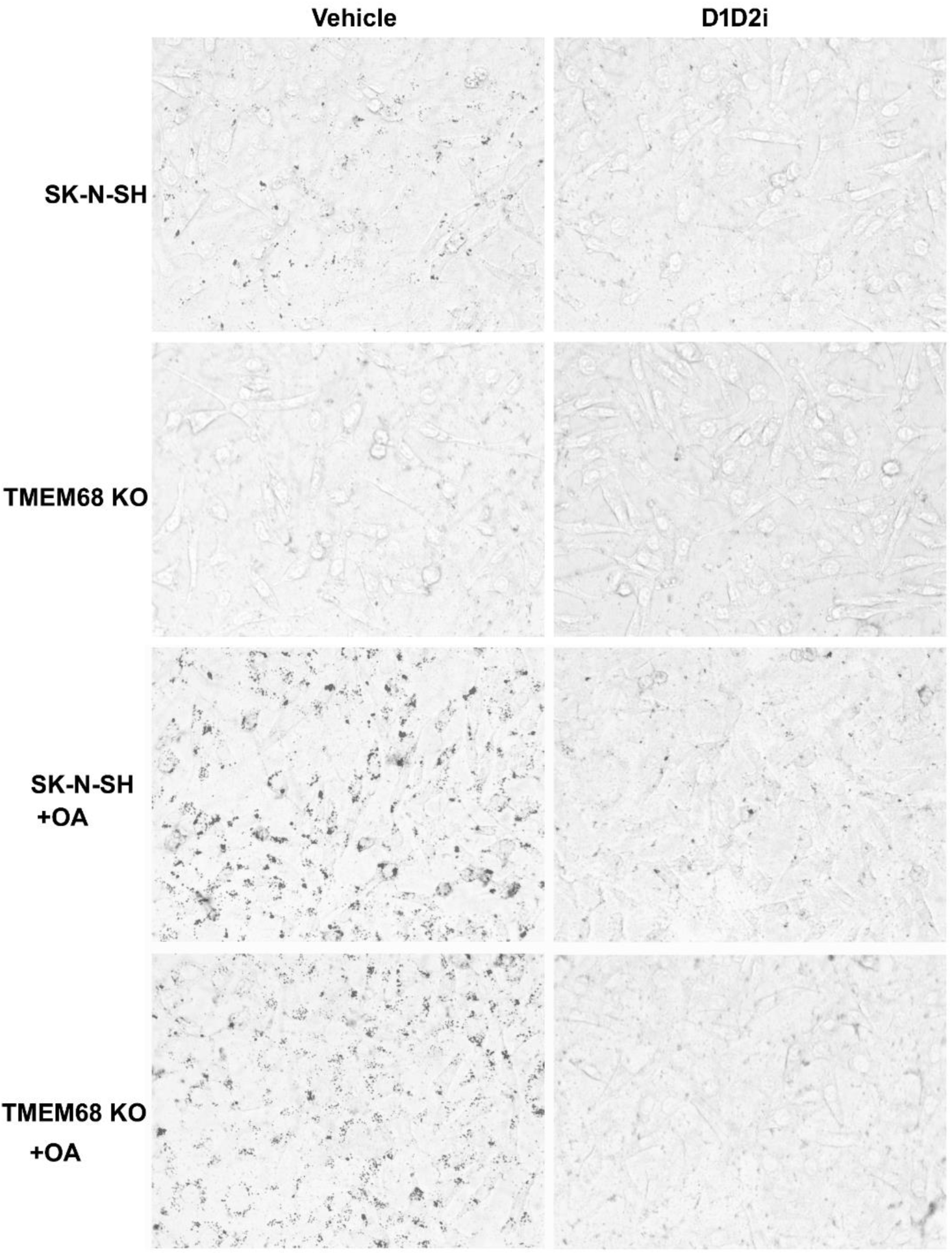
Oil Red O staining of LDs in SK-N-SH and SK/TMEM68 KO cells cultured in the absence or presence of vehicle, a combination of DGAT1 and DGAT2 inhibitors (D1D2i), and OA (+OA) as indicated.

**Figure 4-figure supplement 1.**
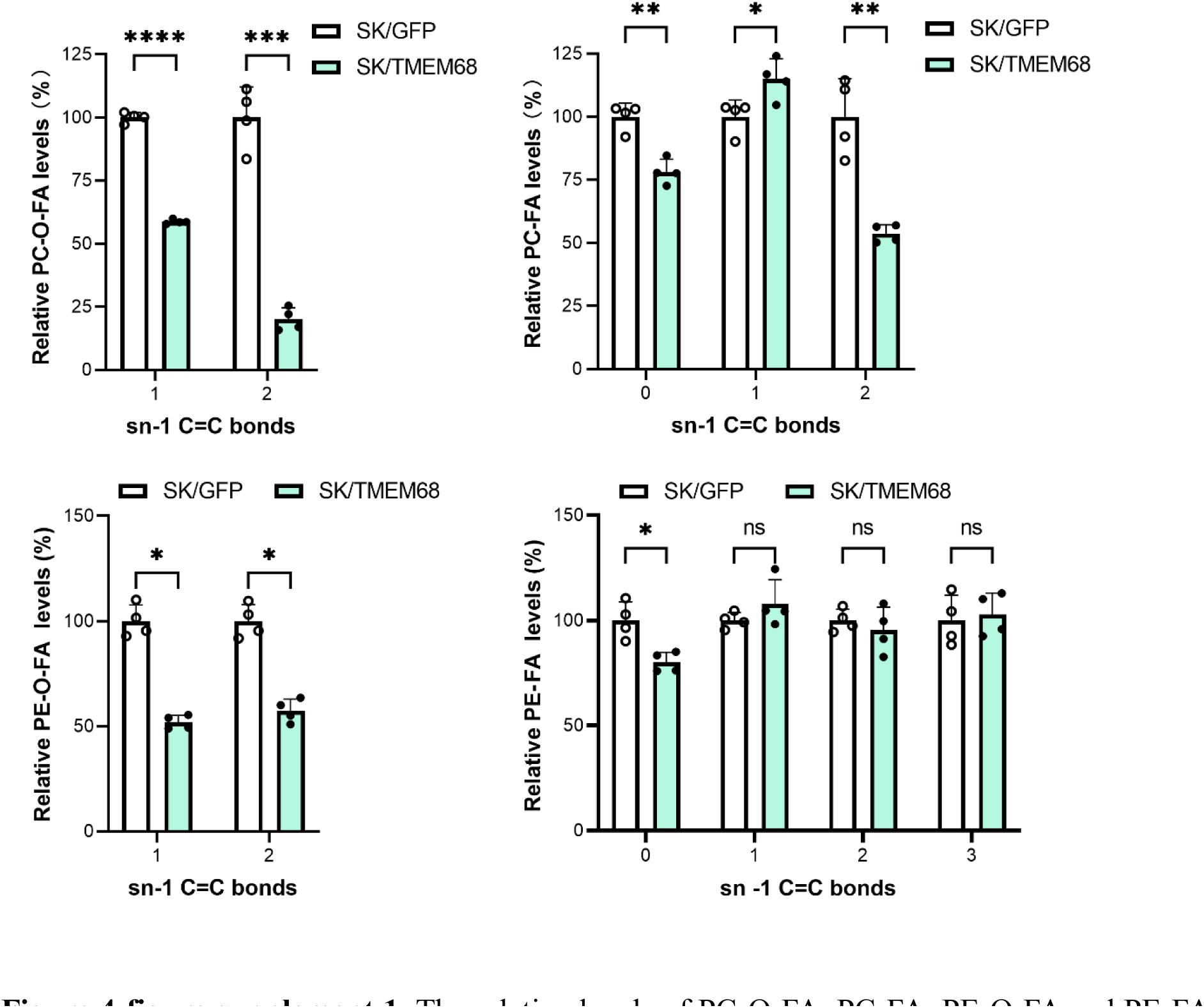
The relative levels of PC-O-FA, PC-FA, PE-O-FA and PE-FA species with different saturation at the sn-1 position (sn-1 C=C bonds) in SK/GFP and SK/TMEM68 cells. Data is presented as means ± SD. Statistically significant differences are marked with asterisks indicating *P*-values. * *P* < 0.05, ** *P* < 0.01, *** *P* < 0.001, **** *P* < 0.0001. ns, not significant. *n* = 4.

**Figure 4-figure supplement 2.**
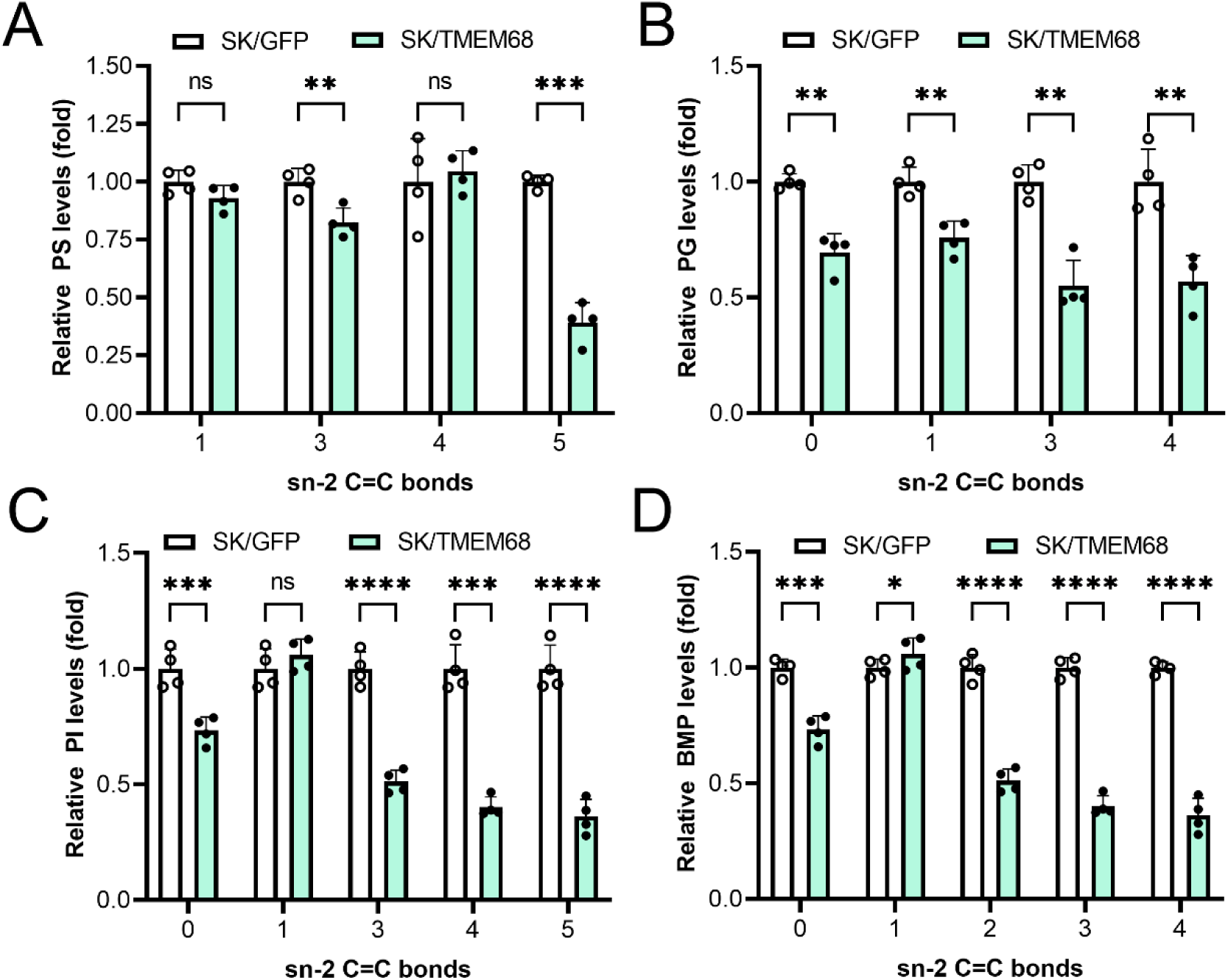
The relative levels of (A) PS, (B) PG, (C) PI and (D) BMP species with different saturation at the sn-2 position (sn-2 C=C bonds) in SK/GFP and SK/TMEM68 cells. Data is presented as means ± SD. Statistically significant differences are marked with asterisks indicating *P*-values. * *P* < 0.05, ** *P* < 0.01, *** *P* < 0.001, **** *P* < 0.0001. ns, not significant. *n* = 4.

**Figure 5-figure supplement 1.**
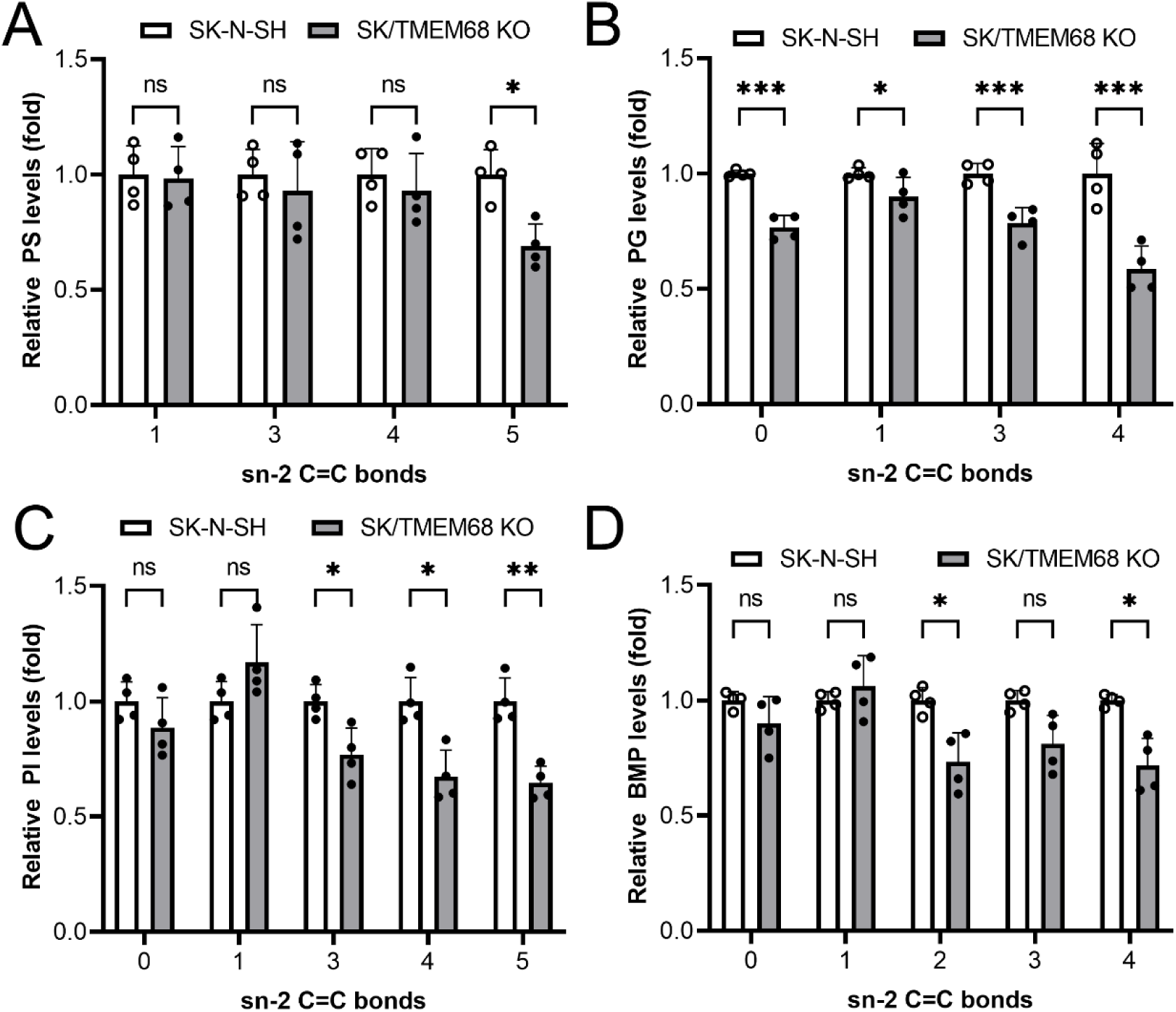
The relative levels of (**A**) PS, (**B**) PG, (**C**) PI and (**D**) BMP species with different saturation at the sn-2 position (sn-2 C=C bonds) in SK-N-SH and SK/TMEM68 KO cells. Data is presented as means ± SD. Statistically significant differences are marked with asterisks indicating *P*-values. * *P* < 0.05, ** *P* < 0.01, *** *P* < 0.001, **** *P* < 0.0001. ns, not significant. *n* = 4.

## Notes

### Competing Interest Statement

The authors have declared no competing interest.

