## Supplementary material for "Human transmembrane protein 68 links triacylglycerol synthesis to membrane lipid homeostasis": Supplementary File 1.docx

**Table supplement 1** Primers used in qPCR experiment

| Primer name | Primer sequence (5' to 3') |
| --- | --- |
| hGPAT3F | ACTTTAGAGTGGGCCACAATACG |
| hGPAT3R | TGGGTGACTCATCTCTTTGGAT |
| hGPAT4F | GGTATCCGCAAACTCTACATGAA |
| hGPAT4R | CCACTTCGACGAATCTCTTTGA |
| hAGPAT3F | CTGCTGGTCGGCTTTGTCTT |
| hAGPAT3R | TCCAGAGTGAGTAGGCGAGG |
| hAGPAT4F | CTCAGGGCTAATCATCAACACC |
| hAGPAT4R | GCTTGAGATGCAATAGGACAGT |
| hATGLF | GAGATGTGCAAGCAGGGATAC |
| hATGLR | CTGCGAGTAATCCTCCGCT |
| hLPIN1F | AGCCTCATACCCTAATTCGGAT |
| hLPIN1R | CCTTTCCGTGGACTTGCTGA |
| hDGAT1F | GGTCCCCAATCACCTCATCTG |
| hDGAT1R | TGCACAGGGATGTTCCAGTTC |
| hDGAT2F | ATTGCTGGCTCATCGCTGT |
| hDGAT2R | GGGAAAGTAGTCTCGAAAGTAGC |
| hFASNF | AAGGACCTGTCTAGGTTTGATGC |
| hFASNR | TGGCTTCATAGGTGACTTCCA |
| hACACAF | TCACACCTGAAGACCTTAAAGCC |
| hACACAR | AGCCCACACTGCTTGTACTG |
| hSCD1F | TTCCTACCTGCAAGTTCTACACC |
| hSCD1R | CCGAGCTTTGTAAGAGCGGT |
| hFABP4F | ACTGGGCCAGGAATTTGACG |
| hFABP4R | CTCGTGGAAGTGACGCCTT |
| hPPARγF | TACTGTCGGTTTCAGAAATGCC |
| hPPARγR | GTCAGCGGACTCTGGATTCAG |
| hGPAT1F | GATGTAAGCACACAAGTGAGGA |
| hGPAT1R | TCCGACTCATTAGGCTTTCTTTC |
| hGPAT2F | TGTGGTCGTCAGGCTTTGG |
| hGPAT2R | GGTCCGTTATGCTTCTGTGGA |
| hCCTαF | GCCCTATGTCAGGGTAACTATGG |
| hCCTαR | GCGTGACCAGAGTGAAATAAGT |
| hCEPT1F | ATGTGGAGATTCTCACCCGGA |
| hCEPT1R | TCTTCTAGCCGCTTTAGTTGGT |
| hCDS1F | GCTTCAGACTTACTCACTTCCAC |
| hCDS1R | GCAAAGGTTGACAGTGCAATG |
| hCDS2F | CACTATTGGCTACAACGTCTACC |
| hCDS2R | AGAAGTAATCCGTCACTGTCTCA |
| hLPCAT1F | CGCCTCACTCGTCCTACTTC |
| hLPCAT1R | TTCCCCAGATCGGGATGTCTC |
| hLPCAT3F | GGAGCTGAGCCTTAACAAGTT |
| hLPCAT3R | CAAAGCAAAGGGGTAACCCAG |
| hLPCAT4F  hLPCAT4R | GGCCTTTATCGTCCTCTTTCTC  CATCCTGTAATTGGCTCCTGAAG |
| hFAR1F | AGACACCACAAGAGCGAGTG |
| hFAR1R | CCAGTTTAGGTTGGGTGAGTTC |
| hFAR2F | CATTGTGGGAGCAACTTGGC |
| hFAR2R | AGTAGCTTTTATGGCCCGAAGA |
| hTMEM68F  hTMEM68R | GCGACCTGCTATGGCAATGA CTCCAACTGCTCCACACCAA |
| hactinF | CATGTACGTTGCTATCCAGGC |
| hactinR | CTCCTTAATGTCACGCACGAT |
