## Supplementary figures and images for "Human transmembrane protein 68 links triacylglycerol synthesis to membrane lipid homeostasis"

### WB source data.pptx

#### Slide 1
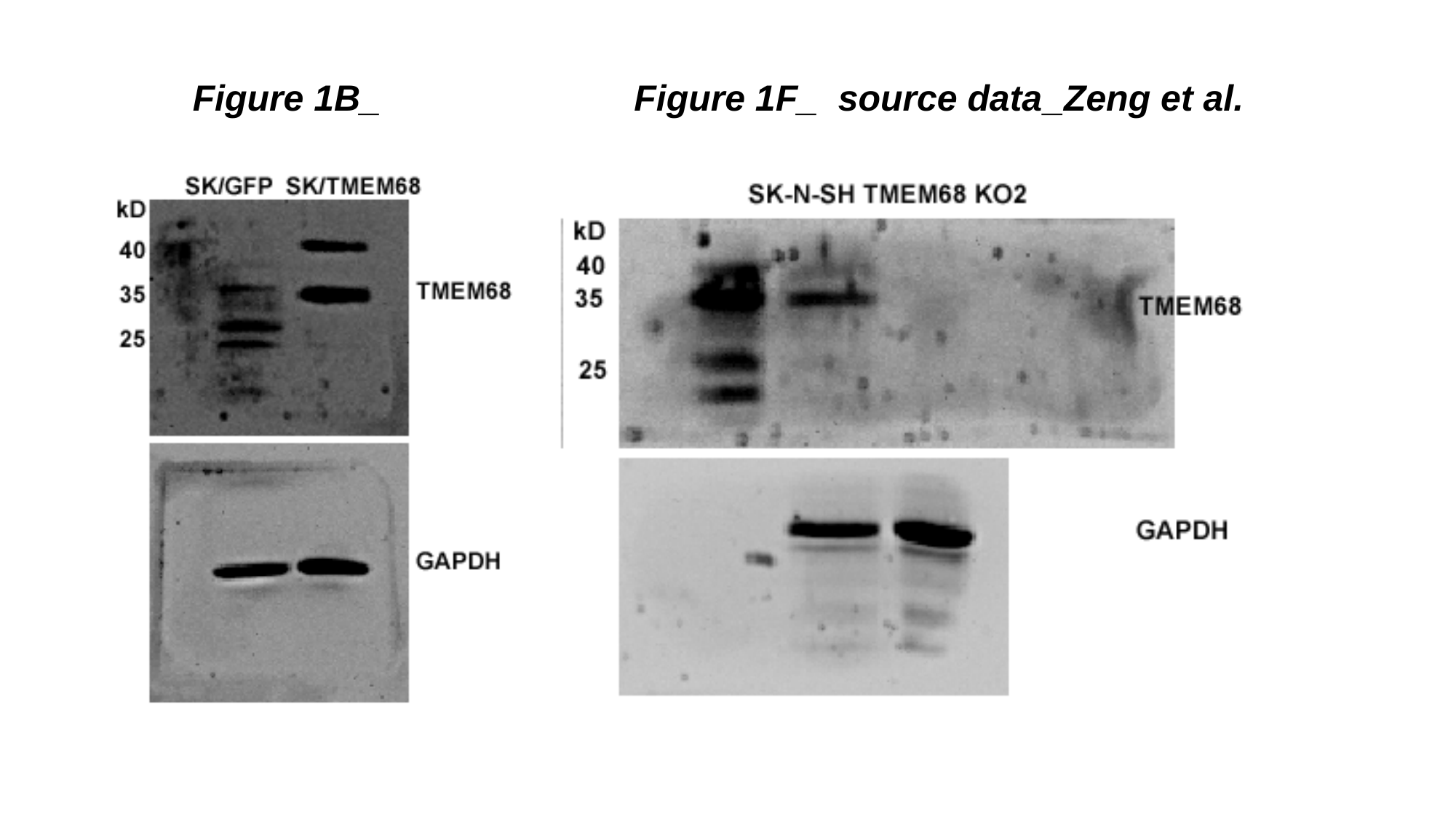

Figure 1B_ Figure 1F_ source data_Zeng et al.
#
